# satmut_utils: a simulation and variant calling package for multiplexed assays of variant effect

**DOI:** 10.1101/2022.04.25.489390

**Authors:** Ian Hoskins, Song Sun, Atina Cote, Frederick P. Roth, Can Cenik

## Abstract

**Background:** The impact of thousands of individual genetic variants on molecular phenotypes for disease-relevant genes remains unknown. Multiplexed assays for variant effect (MAVEs) are highly scalable methods to annotate the relevant variants. However, current software methods for analyzing MAVEs lack standardized annotation, can require cumbersome configuration, and do not easily scale to large target regions.

**Results:** Here, we present satmut_utils as a flexible solution for 1) simulation of saturation mutagenesis data; and 2) quantification of variants across four orders of magnitude from multiplexed assay data. Improvements of satmut_utils over existing solutions include support for multiple experimental strategies, unique molecular identifier-based consensus deduplication, and machine learning-based error correction. We developed a rigorous simulation workflow to validate the performance of satmut_utils and carried out the first benchmarking of existing software for variant calling. Finally, we used satmut_utils to determine the mRNA abundance of thousands of coding variants in cystathionine beta-synthase (*CBS*) by two library preparation methods. We identified an association between variants near chemical cofactor binding sites and decreased mRNA abundance. We also found a correlation between codon optimality and the magnitude of variant effects, emphasizing the potential of single-nucleotide variants to alter mRNA abundance.

**Conclusions:** satmut_utils enables high-performance analysis of saturation mutagenesis data, achieves unprecedented specificity through novel error correction approaches, and reveals the capability of single-codon variants to alter mRNA abundance in native coding sequences.

## Background

Multiplexed assays of variant effect (MAVEs) employ next-generation sequencing to profile the phenotypic effects of hundreds to thousands of genetic variants in a target gene. These assays, which rely on saturation mutagenesis, have been used to survey variant effects on molecular phenotypes ranging from mRNA and protein expression to protein binding and enzyme activity [1–10]. As a result, MAVEs emerged as methods to study variant effects and annotate variant significance, ultimately informing disease diagnosis and prognosis [11, 12]. Guidelines now exist for the development of MAVEs, underscoring their utility for variant annotation and interpretation [13]. Given saturation mutagenesis data contains variants with frequencies at and even below error rates for some polymerases (1 x 10^-4^), variant callers for MAVEs must have not only high sensitivity, but also high specificity. Yet, analysis methods for MAVEs are not standardized, and to our knowledge, none of the existing variant callers for analysis have previously benchmarked performance.

Existing tools for MAVE analysis require detailed configuration of parameters (**Methods**), and may only address particular experimental designs. For example, the dms_tools2 package [14] has specific input requirements: primer designs should start flush with codons, and reads must be dual-barcoded and align contiguously to a user-provided reference (no insertions or deletions). Similarly, the Enrich2 package [15], requires that reads align contiguously with a provided reference sequence. DiMSum [16] only annotates standard amino acid changes, and like dms_tools2 and Enrich2, only calls variants in a single PCR amplicon at a time. Hence, none of the current methods allow variant calling from multiple PCR amplicons at once with one configuration of analysis parameters, limiting the ability to rapidly scale to large genes. Also, current strategies generally assume pre- and post-selection sequencing of the variant library, for example when assaying variant effects on organismal growth [4]. While this is the predominant MAVE design, a generalized variant caller would facilitate not only selection-based assays but also assays of arbitrary design.

Similarly, while a multitude of methods exist to call somatic variants in clinical samples [17–19], somatic variant callers for whole-transcriptome analysis are tailored to quantify variants in samples with few real <u>s</u>ingle- and <u>m</u>ulti-nucleotide polymorphisms (<u>SNP</u>s and <u>MNP</u>s, respectively). In contrast, MAVE data contain a high density of low-frequency SNPs and MNPs. For example, di- and tri-nt MNPs may comprise a large proportion of the total variants in codon saturation mutagenesis. The low frequency of variants (< 1 x 10^-4^) pose new problems to variant calling for MAVE data. Analysis is further complicated by the hierarchical composition of variants, wherein true positive variants may be called together with nearby true or false positive variants [16].

To address the need to call ultra-low frequency variants in MAVE data, we designed and implemented satmut_utils (<u>sat</u>uration <u>mut</u>agenesis <u>util</u>itie<u>s</u>), incorporating modern software development practices, extensive documentation, integration with package management, and rigorous unit testing. The satmut_utils ‘call’ workflow is an end-to-end variant caller for MAVEs that supports analysis of both a) amplicon [4]; and b) rapid-amplification-of-cDNA-ends (RACE)-like library preparation methods [20–22]. To achieve high specificity, satmut_utils optionally builds on a simulation workflow (‘sim’), enabling the generation of datasets for benchmarking and error modeling.

Here, we performed the first benchmarking analysis of MAVE variant callers and show that satmut_utils achieves superior performance for MAVE analysis. We then assayed variant effects on mRNA abundance using two library preparation methods. Using satmut_utils, we identified variants in cystathionine beta-synthase (*CBS)* with effects on mRNA abundance, expanding a prior variant effect map for *CBS* function [10]. We further characterized possible mechanisms of altered mRNA abundance, including codon-mediated, *cis-*regulatory effects. The satmut_utils package enables flexible experimental design and comparative analysis of saturation mutagenesis data from various sources, and will facilitate the interpretation of variant effects on the RNA life cycle.

## Results

### Design of simulation and variant calling workflows for saturation mutagenesis data

We developed a workflow to simulate ultra-low frequency variants in real alignments, termed ‘sim’ (**Figure 1A**). ‘sim’ generates variants by editing into pre-existing alignments that correspond to a negative control (NC) sequencing library prepared from a non-mutagenized template. Editing real alignments enables us to capture sequencing errors and experiment-specific biases that may escape model-based *in silico* read generation. ‘sim’ can efficiently simulate the number of variants typically targeted in MAVEs (>1000) in a single transcript, improving on the scalability of existing solutions [23].

**Figure 1:**
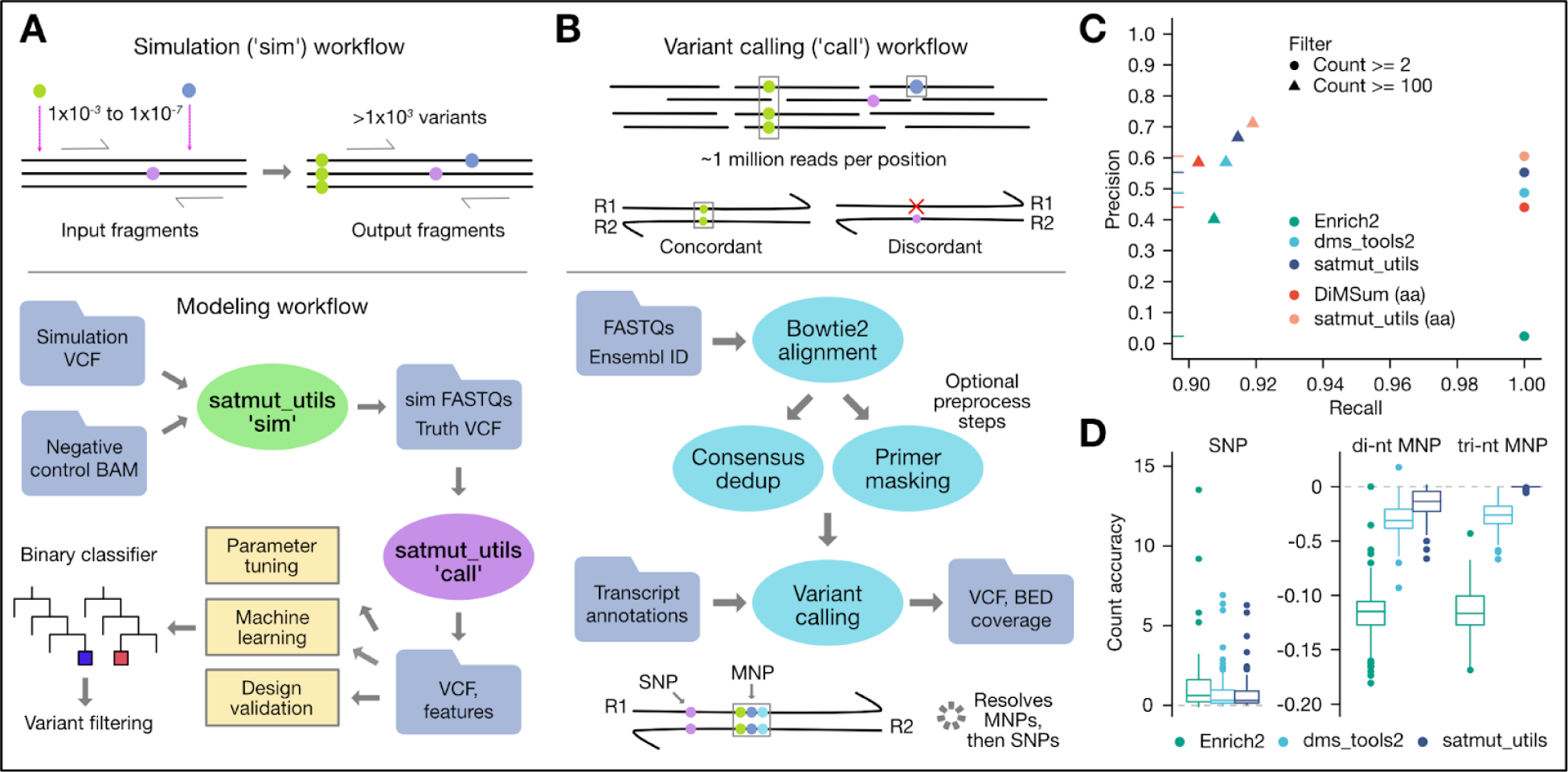
satmut_utils design and performance benchmarking. Solid circles represent single or multiple-nucleotide polymorphisms (SNPs, MNPs), which may be either true or false positives (errors). **A) Variant simulation workflow.** With ’sim’, ultra-low frequency variants in Variant Call Format (VCF) are edited into pre-existing sequencing read alignments (BAM). Edited reads (FASTQ) and true positive variants (Truth) are output with expected counts and frequencies. The ’call’ workflow (**B**) extracts quality features during variant calling, which may be used for assay design validation, software parameter tuning, and machine learning-based error correction. **B) Variant calling workflow**. SNPs and MNPs are identified and quantified in paired-end reads following optional preprocessing to improve specificity. Transcript nucleotide and protein changes are annotated and a VCF and fragment coverage bedgraph file are output. **C) Performance of MAVE variant callers.** 281 variants were simulated in alignments for a single amplicon in *CBS* and performance measures were evaluated after applying to simple count filters. nt: nucleotide/codon-level calls; aa: amino acid-level calls. **D) Accuracy of variant count estimates.** Count accuracy is quantified as the difference between the observed and truth (simulated) count, divided by the truth count. One outlier SNP was excluded for visualization (Enrich2 accuracy: 130.6; satmut_utils and dms_tools2 accuracy: 1.415). dedup=deduplication.

The ‘sim’ workflow supports editing of multiple SNPs and MNPs at the same coordinate; ensures reads are edited only once; and allows the user to prohibit simulation of variants adjacent to pre-existing errors to ensure errors in the edited read do not convert the simulated variant to higher order (e.g. SNP to MNP; see **Additional File 1**). In summary, satmut_utils ‘sim’ enables deterministic simulation of many low frequency variants at the same position, and offers the first generalized simulation method specific for multiplexed assays.

Next, to call ultra-low frequency SNPs and MNPs in targeted sequencing data, we developed the satmut_utils ‘call’ workflow (**Figure 1B**). Importantly, ‘call’ supports variant calling from multiple interleaved PCR tiles simultaneously, a feature lacking from other tools for analysis [14,15,24]. A curated human transcriptome is included to facilitate ease-of-use, although custom reference files are also supported. Our method provides two additional features missing in other MAVE analysis methods. First, satmut_utils enables variant calling from RACE-like library preparation methods such as Anchored Multiplex PCR [20] (**Supplementary Figure 1A**). Second, satmut_utils extracts read-based quality data for each mismatch contributing to a primary variant call. Quality data may then be used to train error correction models. (For a detailed comparison of variant caller features see **Supplementary Figure 1B**; for time and memory consumption of satmut_utils see the **Additional File 1**).

To improve specificity of variant calls, the ‘call’ algorithm (**Supplementary Figure 1C**) incorporates filters based on read edit distance and base qualities. Then, variants are called in read pairs if mates are concordant [4, 10], i.e. if the same base call is observed in both forward and reverse reads. This filters out sequencing errors which are found in only one read of the pair. Finally, satmut_utils employs a novel variant calling algorithm that prioritizes MNPs and improves sensitivity for MNP calls when they are adjacent to errors (**Supplementary Figure 1D**). We coined the term *variant conversion* for cases when a true variant and adjacent error are called together as a false positive (**Additional File 1**). Conversion is particularly insidious for MAVE analyses as it may also lead to a false negative call. Altogether, satmut_utils 1) requires a single configuration for analysis of data from multiple amplicons; 2) supports two different library preparation methods; and 3) employs a unique variant calling algorithm for high-accuracy estimates of variant abundance.

### *In silico* validation and benchmarking of variant calls with ‘sim’

We compared performance of satmut_utils to dms_tools2 [14], Enrich2 [24], and DiMSum [16], in the first benchmarking analysis of MAVE variant callers. We speculate that a prior lack of benchmarking was due to several challenges: 1) lack of truth datasets; 2) different experimental design assumptions; and 3) non-standardized input and output file formats. Nonetheless, after preprocessing alignments to meet the various input requirements for Enrich2 and dms_tools2 (**Methods**), we successfully generated a common benchmarking dataset using reads from a single PCR amplicon in cystathionine beta-synthase (*CBS)* [10]. This simulated dataset contained 281 variants at frequencies between 1 x 10^-6^ to 1 x 10^-3^ in a background of approximately two million negative control read pairs.

With a threshold of two supporting reads/fragments to make a variant call, Enrich2, dms_tools2, and satmut_utils achieved perfect sensitivity at the nucleotide level (**Figure 1C**). However, precision was 0.023 (Enrich2), 0.487 (dms_tools2), and 0.553 (satmut_utils). Because DiMSum does not output annotations at nucleotide resolution, we compared DiMSum to satmut_utils for amino acid changes. At perfect sensitivity, DiMSum precision was 0.440 compared to satmut_utils precision of 0.605. Lower precision for Enrich2 and DiMSum may be due to merging with nearby errors. The satmut_utils ‘call’ workflow does not call phased SNPs as a MNP unless the SNPs are within 3 nt (no haplotype calls are made). We found that this algorithmic design choice is a reasonable compromise to remove thousands of false positive calls arising from the merging of read errors. We note that Enrich2 precision might be higher with another analysis mode (barcoded sequencing). See **Additional File 1** for a detailed explanation of benchmarking considerations.

Despite differences in overall performance, dms_tools2 and Enrich2 reported largely similar counts to satmut_utils for true positive variants, especially MNPs (**Supplementary Figure 2B,C**). Yet, strikingly, satmut_utils reported more accurate variant counts than other methods for MNPs (**Figure 1D**). Deviations from the truth count are likely impacted by read filtering and the variant calling algorithm (**Supplementary Figure 1 C,D**), which may explain the higher accuracy of satmut_utils variant calling. In total, by satmut_utils ‘sim’, we performed the first benchmarking analysis of MAVE variant callers, and showed that satmut_utils ‘call’ is more accurate than other methods for variant calling of *in silico* mutagenesis data.

### ‘sim’ and ‘call’ power machine learning-based error correction

Sequencing libraries contain systematic errors arising from library preparation- and sequencer-specific biases [25–27]. In MAVEs, a negative control (<u>NC</u>) library of the non-mutagenized template is typically sequenced in the same experiment as mutagenized libraries [10]. In agreement with prior observations of experiment- and platform-specific errors [27], we found a wide range of error rates for independent libraries from various labs, experiments, and sequencing runs (**Figure 2A**). We noted the highest error rates for (C>A, G>T) and (C>T, G>A) substitutions across all Illumina platforms, library preparation methods, and independent libraries from various input nucleic acid sources. We hypothesized that sequencing a NC library, simulating variants in this control, and then training classifiers would help moderate such biases.

**Figure 2:**
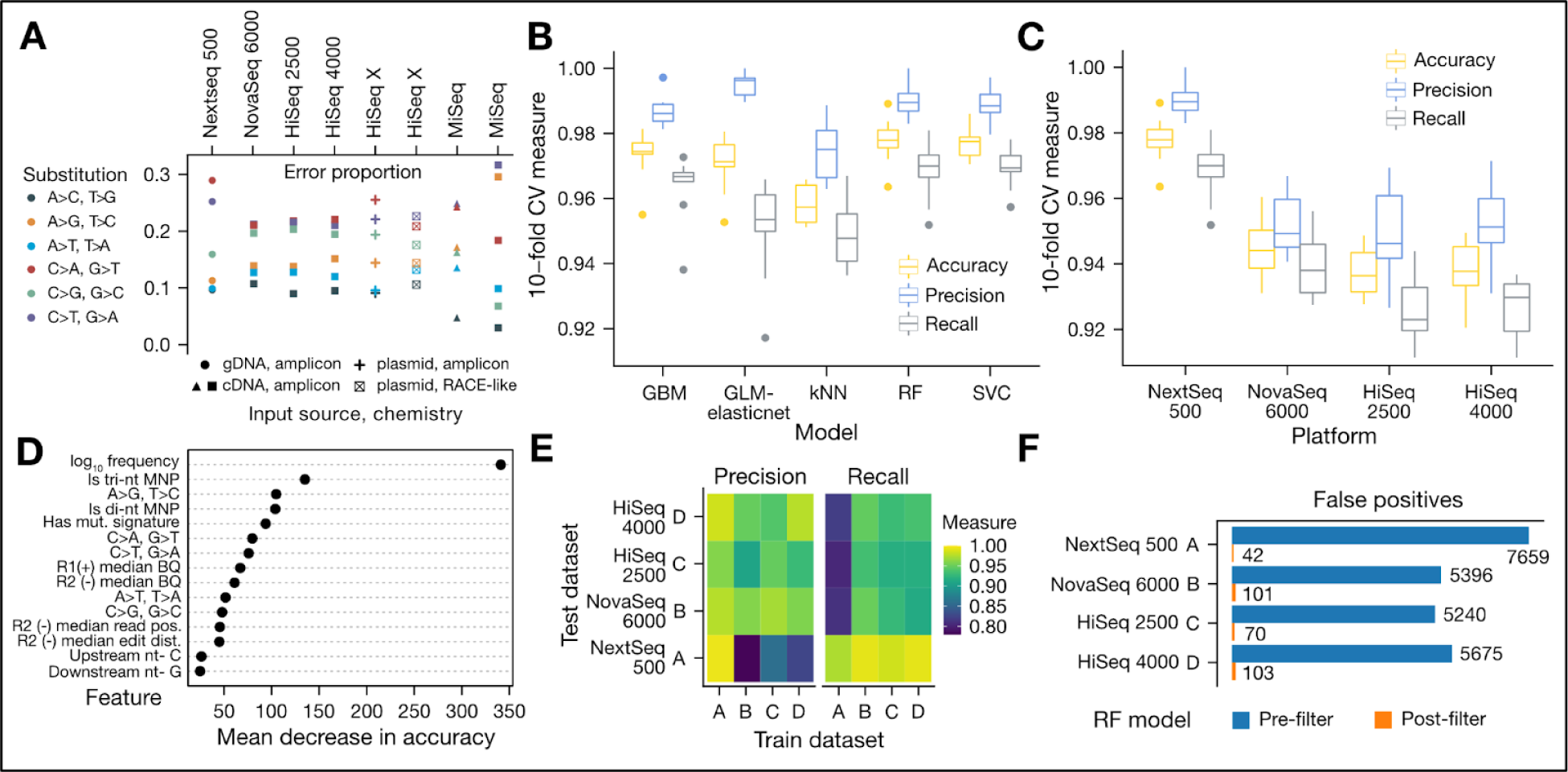
Machine learning models for error correction. Negative control (NC) alignments for **‘**sim’ dataset *A* (Nextseq 500) arose from the human *CBS* coding sequence after functional complementation in yeast [10]. Alignments for ‘sim’ datasets *B-D* (NovaSeq 6000, HiSeq 2500, HiSeq 4000) and MiSeq runs arose from HEK293T endogenous *CBS* cDNA, and alignments for HiSeq X datasets arose from *CBS* template plasmid. **A**) **Error proportions in negative control libraries**. Proportion of each error substitution across NC libraries from various sources. Shape of the points indicates an independent NC library. **B) Model selection.** To compare models, dataset *A* (3802 variants, 7859 true mismatches, 6463 false mismatches) was used. Up to 19 satmut_utils call quality features were selected to train binary classifiers (Methods). **C) Random forest performance.** Random forests (RF) were trained on all four ‘sim’ datasets and cross-validation performance across different platforms was calculated. **D) Feature importance for RF models.** A RF was trained on a combined dataset (all ‘sim’ datasets *A-D*), and the top fifteen important features as measured by mean decrease in accuracy (Methods) are plotted. **E) Cross-generalization of RF models.** Pairwise train-test regimes were carried out with all ‘sim’ datasets to assess model generalization across sequencing libraries and platforms. **F) Error correction impact on variant calls in NC libraries.** satmut_utils variant calls from each NC library were filtered by the RF models. The number of error mismatches before and after filtering is plotted for each NC library. NC: negative control; GBM: gradient boosted machine; GLM-elasticnet: generalized linear model with elastic net regularization; kNN: k-nearest neighbors; RF: random forest; SVC: support vector classifier.

To test the utility of error correction models enabled by satmut_utils, we generated four large simulated datasets by editing thousands of variants into two NC libraries, sequenced on four Illumina platforms (**Methods**). At perfect recall, satmut_utils precision in these datasets with default calling parameters and no model-based error correction (**Methods**) was 0.552 +/- 0.041 (mean +/ s.d.). Thus, with a naïve filter using a minimum count threshold, thousands of false positives remain in multiplexed assay datasets, deteriorating their quality.

We next used the simulated dataset from the first NC library, which comprised the human *CBS* coding sequence after functional complementation in yeast [10], to assess performance of machine learning models in reducing false positives. We trained binary classifiers using quality features extracted by satmut_utils ‘call’ from the first simulated dataset (hereafter dataset *A*). Of the five classifiers tested by nested cross validation (CV), all five models showed a median accuracy >0.95 (**Figure 2B**). The remaining three datasets (*B-D*) arose from a second NC library consisting of the *CBS* coding sequence amplified from human HEK293T total RNA and sequenced on different platforms. We selected the random forest (RF) to test performance on all four datasets generated with ‘sim’ (**Figure 2C**). The mean accuracy of the final models (N=4) on an independent test set was 0.954 +/- 0.020 (mean, s.d.), indicating that models trained on simulated data are robust to different choices of NC library and sequencing platform, and outperform filtering variants using a fixed count threshold (Figure 1D). Several quality features lent predictive power as measured by RF feature importance (**Figure 2D**).

To assess generalization of the models, we trained a RF on one simulated dataset and tested it on all other datasets (all pairwise permutations, **Figure 2E**). Models generalized well for our own NC library sequenced on different platforms, with an accuracy of 0.939 +/- 0.014 (mean, s.d.). Accuracy was slightly worse when trained on the independent dataset *A* and tested on datasets *B-D*: 0.892 +/- 0.010 (mean, s.d.). We finally applied the models to filter calls in the NC libraries and observed a strong reduction of false positives (**Figure 2F**). Therefore, training error correction models on simulated data significantly improves variant calling precision (see **Additional File 1** for potential caveats).

### Read pre-processing steps implemented in satmut_utils reduce false positive variant calls

While machine learning models using sequence-level features significantly reduce false positives, additional improvements that leverage read pre-processing can further improve specificity. For example, primer base quality masking [19] may be used to omit variant calls that have arisen from primer synthesis errors by setting base qualities to 0 for synthetic read segments (**Figure 3A**). When unique molecular indices (UMIs) are incorporated into the library design, further improvements can be obtained by consensus deduplication [14,19,28], where a consensus sequence is generated from PCR duplicates (**Figure 3B**). We implemented these additional methods and compared variant calls in simulated datasets before and after primer masking and consensus deduplication using UMIs.

**Figure 3:**
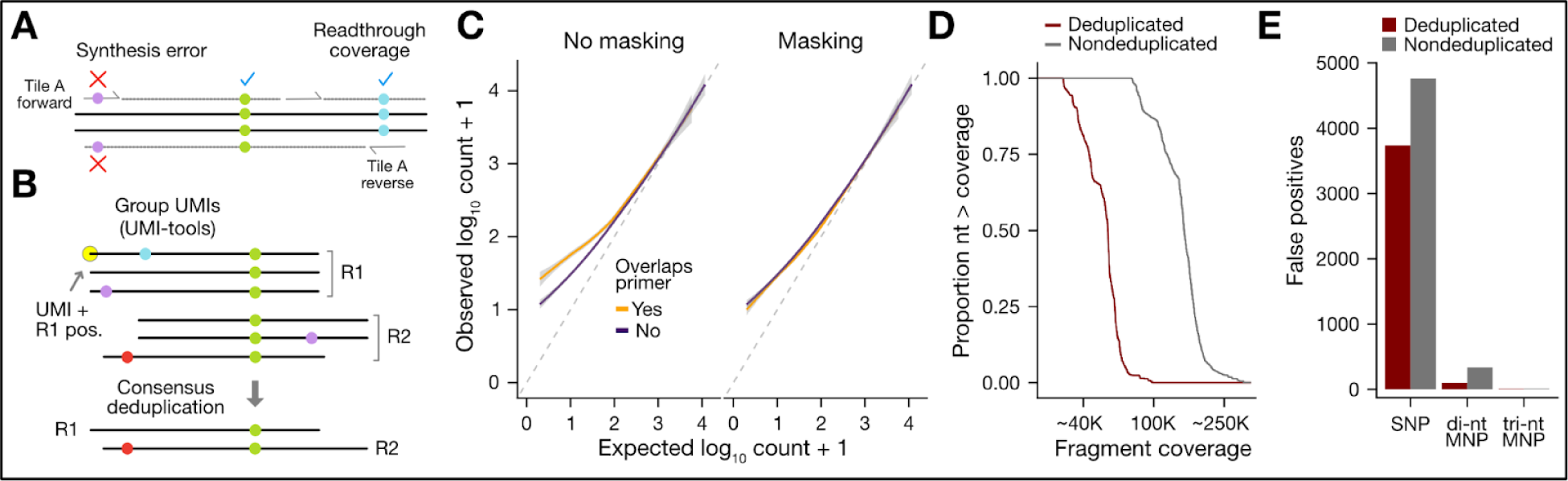
Read preprocessing for error correction. For A and B, solid colored circles represent SNPs, either true or false positive (error). **A) Primer base quality masking schematic.** Base qualities for read segments determined to originate from synthetic primer sequences are set to 0. Black lines indicate the sequenced fragment. Solid gray lines with ticks represent primers/directionality. Grey dotted lines represent reads off the input fragment. Readthrough coverage refers to coverage from adjacent PCR tiles, required to call variants that overlap primers. **B) Consensus deduplication schematic.** UMI-tools directional adjacency method [73] is used to group paired-end reads from a common unique fragment, defined by UMI and read 1 position (R1 pos.). A custom consensus deduplication algorithm generates the consensus base among duplicates at each aligned fragment position for each read. **C) Primer base quality masking improves accuracy of variants underlying primers.** Simulated datasets (Figure 2, N=4) were analyzed with/without primer BQ masking and true positive variants that overlap primers are plotted compared to variants not overlapping a primer. **D) Consensus deduplication maintains coverage uniformity.** A UMI-containing, RACE-like negative control (NC) library was generated. Waterfall plots of cumulative fragment coverage for consensus-deduplicated reads and non-deduplicated reads indicate uniform collapse of PCR duplicates. x-axis is in log_10_ scale with a range between 4.6 and 5.6. **E) Consensus deduplication reduces false positives.** The effect of consensus deduplication is shown for the RACE-like NC library for each variant type. UMI=Unique molecular index; SNP=Single nucleotide polymorphism; MNP=Multiple nucleotide polymorphism; NC: negative control.

Primer masking removed a small number of false positive SNPs in ‘sim’ datasets (min=1, max=64; min proportion of SNPs remaining=0.985). More importantly, primer base quality masking improved the accuracy of variant counts in simulated data (**Figure 3C**, N=4 pooled datasets). In parallel with primer masking, consensus deduplication of a RACE-like (Anchored Multiplex PCR) NC library through UMIs reduced depth of coverage across *CBS* by 63.1% (**Figure 3D, Methods**). Further, deduplication reduced false positive (FP) SNPs by 21.5% (1026 FPs); di-nt MNPs by 70.3% (237 FPs); and tri-nt MNPs by 27.2% (3 FPs) (**Figure 3E**). This significant improvement in specificity may be accompanied by a slight cost to sensitivity, but the current implementation of the ‘sim’ workflow was insufficient to determine the exact sensitivity-specificity tradeoff (see **Additional File 1** for details). Altogether, read pre-processing steps can improve the quality of MAVE data prior to variant calling, independent of other model-based error correction.

### Variant calls are reproducible by two orthogonal library preparation methods

To demonstrate the flexibility of satmut_utils in analyzing MAVE data, we measured gDNA and mRNA abundance for a complete coding variant library in *CBS* following stable expression in a HEK293T landing pad cell line (iCasp9 Int Blast) [29]. We recombined a *CBS* variant library [10] into the landing pad line with a downstream IRES-mCherry element (**Figure 4A**). Then, we assayed variant abundance in gDNA and cDNA by amplicon [4, 10] and RACE-like (Anchored Multiplex PCR) [20] library preparation methods (**Methods**). The quality of total RNA input as well as PCR products at steps in library preparation were confirmed (**Supplementary Figure 3A-C**).

**Figure 4:**
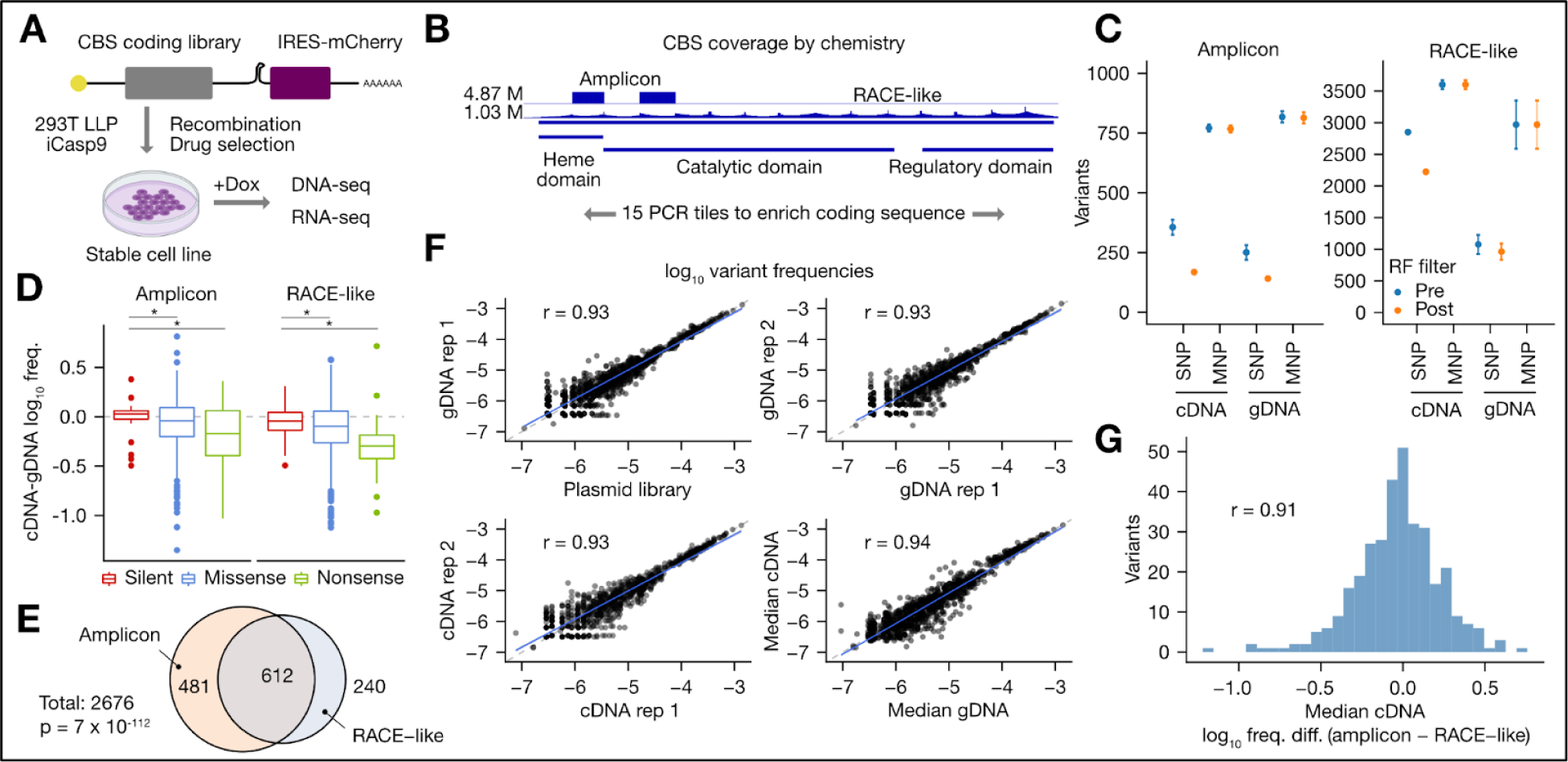
satmut_utils analysis of a *CBS* variant library by orthogonal library preparation methods. **A) Experimental strategy.** A human *CBS* coding variant library was stably expressed in a landing pad cell line (293T LLP iCasp9 Blast) [29]. gDNA and total RNA were sequenced by one of two targeted-sequencing methods 24 h after induction with Doxycycline (Dox). **B) *CBS* domains and sequencing coverage.** Coverage for two tiles by the amplicon method is contrasted with full coverage in the RACE-like method. The maximum coverage depth is shown on the left of the track. **C) Filtering of variant calls.** Random forest models were trained for each method using negative control libraries and the ‘sim’ workflow. Plotted are the mean number of variant calls +/- the standard deviation (N=3). The total possible calls are as follows: amplicon (531 SNPs, 2145 MNPs); RACE-like (4105 SNPs, 16863 MNPs). **D) Differential abundance for mutation types.** The difference in the median log_10_ frequency between cDNA and gDNA is shown for variants observed in all gDNA and cDNA replicates (N=6). Outliers are greater than 1.5 * interquartile range. Asterisks indicate significant differences by one-sided Wilcoxon rank sum tests (p < 0.05). **E) Variant call overlap between methods.** Overlap in variant calls is shown for *CBS* tiles 2 and 4. Total calls is the theoretical number of variant calls. Significance of overlap was computed by a hypergeometric test. **F) Biological replicate correlation for amplicon libraries**. Reproducibility between log_10_ frequencies was determined by Pearson’s correlation coefficient, after filtering out variants only observed in one replicate (Methods). **G) Similarity of variant frequency estimates between methods in cDNA replicates.** The difference in median log_10_ frequency between amplicon and RACE-like methods is shown for cDNA libraries with filtering as in F. Reproducibility of cDNA frequencies between methods was determined by Pearson’s correlation coefficient after filters (Methods). RACE=rapid amplification of cDNA ends; M=million; PLP=pyridoxal-5’-phosphate; SNP=single nucleotide polymorphism; MNP=multiple nucleotide polymorphism.

For each method, we included a NC sequencing library to enable variant filtering with a random forest (RF) model. These libraries showed uniform coverage across the CBS target regions (**Figure 4B**), as did mutagenized libraries. We found high performance of RF models for both library preparation methods (0.975, 0.959 accuracy for amplicon and RACE-like simulated datasets, respectively). These models were subsequently used to filter variant calls from the mutagenized sequencing libraries (**Figure 4C**).

The difference in log_10_ frequencies between cDNA and gDNA for each variant highlighted large effects on relative abundance in both directions. As expected, missense and nonsense variants reduced mRNA abundance compared to silent changes (**Figure 4D**). Variant calls made by both library preparation methods comprised 22.9% of the maximum theoretical calls for the two amplicons (**Methods**), and overlap was significant by a hypergeometric test (**Figure 4E**, p=7 x 10^-112^). For the amplicon method, among variants detected at least once, 55.7% of variants were found in all replicate gDNA and cDNA libraries (N=3 replicates each). In contrast, only 15.9% of variants were observed in all replicates by the RACE-like method.

Variant abundance estimates were reproducible across input sources and independent biological replicate cell lines (**Figure 4F**, **Supplementary Figure 4A-B**). Despite a difference in coverage depth (**Figure 4B**), the variant frequency correlation was satisfactory compared to the RACE-like method for variants that were well-measured (0.91 Pearson’s correlation, **Figure 4G**, **Supplementary Figure 4C**). Taken together, our results suggest satmut_utils reports reproducible variant frequency estimates from two library preparation strategies, and facilitates analysis of data from multiple nucleic acid sources.

### Identification of *CBS* variants that alter mRNA abundance

To apply satmut_utils variant calling to unveil biological insights, we next determined *CBS* variants with effects on mRNA abundance. The human CBS enzyme has specific amino acids that bind to two cofactors (heme and pyridoxal phosphate- PLP) [30–32]. These cofactors regulate folding, stability, and activity of CBS [10,33–35]. Because heme and PLP can stabilize CBS variants and remediate pathogenic phenotypes [10,36,37], and because heme binding is not reversible [38], we hypothesized heme facilitates co-translational folding of CBS, similar to its role in folding of globin [39, 40].

We reasoned that *CBS* variants with low mRNA abundance may be enriched at or near important structural residues of CBS, as improper co-translational folding may trigger ribosome quality control, leading to mRNA and protein degradation [41, 42]. To address this hypothesis, we determined *CBS* variants with significant differential abundance between cDNA (total RNA) and gDNA using the high-quality data from the amplicon method (**Figure 5A**, **Supplementary Figure 5A, Methods**). Of 2676 theoretical variants (SNPs, MNPs) for the amplicon method, 1238 were detected (46.3%) at least once, and 691 were observed in all gDNA and cDNA replicates (N=6, 25.8%). Of these, 6 variants were higher in mRNA abundance compared to 22 variants that were lower (FDR < 0.1).

**Figure 5:**
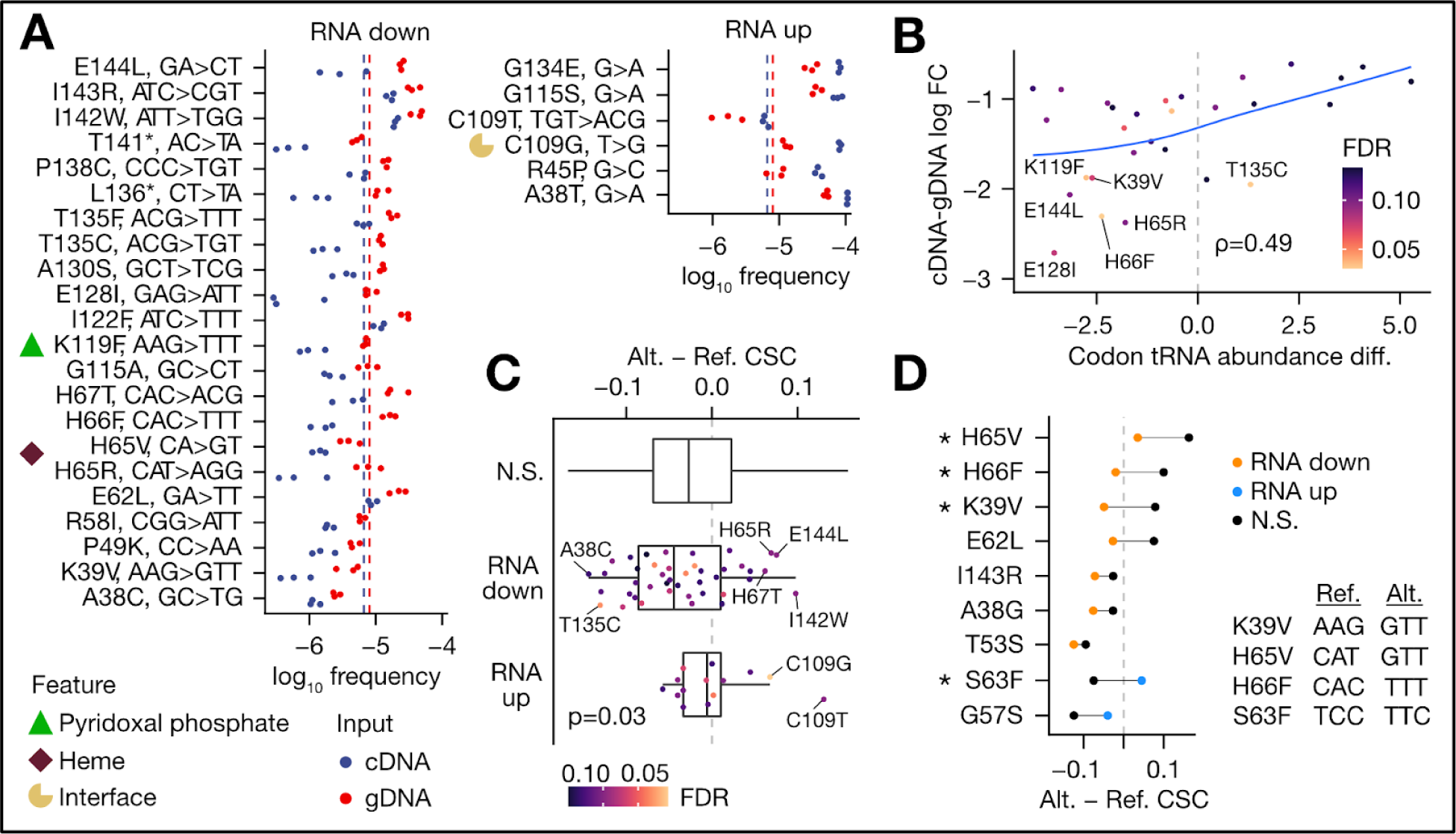
Identification and mechanisms of CBS variants that alter mRNA abundance. In all panels, data from the amplicon method is shown, and a grey dotted line indicates no change in variant effect or comparative metrics. For panel A, variants at a FDR < 0.1 are shown; in all other panels, variants with a FDR < 0.15 were analyzed. **A) CBS variant differential abundance.** y-axis labels denote the amino acid change and the nucleotide substitutions for variants with significant differential abundance. Structural residues at or adjacent to important features are labeled with an icon. Red and blue dotted lines represent the median for each input source. **B) tRNA abundance correlation with mRNA abundance effects.** For each variant, the difference in tRNA abundance (Alt. codon - Ref. codon; hydro-tRNAseq [54]) is plotted against the log fold change. Spearman rank correlation (rho) is shown. Blue line is a loess fit with span=2; confidence intervals were omitted for clarity. **C) Codon stability coefficient for variants grouped by directional effect.** The difference in HEK293T ORFome codon stability coefficient (CSC) scores [50] between alternate and reference codons is compared. p-value indicates a one-sided Wilcoxon rank-sum test between RNA down and RNA up groups. **D) Comparison of codon stability between significant and non-significant variants**. Significant variants were compared to another variant leading to the same amino acid change. Asterisks indicate a difference in the upper quartile of the distribution of all pairwise distances between significant and non-significant variants (null distribution, Methods). PLP=pyridoxal-5’-phosphate; FC=fold change; Ref.=reference codon; Alt.=alternate codon; N.S.=not significant; CSC=codon stability coefficient.

Several variants at and near important residues for activity, including variants at positions previously implicated in CBS deficiency, exhibited significant effects on mRNA abundance (**Supplementary Table 1**). Specifically, we identified decreased mRNA abundance of the variant K119F, which interrupts the Schiff base formed by this residue with PLP [30]. Similarly, mutations at or adjacent to the heme binding residue (H65) exhibited a strong reduction in mRNA abundance: H65R, H65V, H66F, and H67T. Variants with decreased mRNA at positions previously implicated in CBS deficiency were P49K, R58I, E128I, I143R, and E144L [35,43–46].

Twenty other variants had differential mRNA abundance by the RACE-like method (**Supplementary Figure 5B**, FDR < 0.1), and we found modest correlation in the measurement of mRNA abundance effect between amplicon and RACE-like methods (0.57 Pearson’s coefficient, **Supplementary Figure 5C**). We noted variant effects that depend on the position of the variant in the coding sequence. In the amplicon method, the variance of the effect at each position was higher in the catalytic domain than in the heme domain, suggesting the magnitude of *CBS* variant effects may depend on the encompassing structure of the CBS protein (Levene’s test, p=0.005, **Supplementary Figure 6A**). Similarly, RACE-like data indicated nonsense variants had the strongest effects on mRNA abundance when located near the middle (catalytic domain) of the coding sequence (**Supplementary Figure 6B**).

Altogether, we identified *CBS* variants near important functional residues that alter mRNA abundance. Consistent with co-translational folding of CBS by cofactors, mutations at and adjacent to the heme and PLP binding residues uniformly exhibited decreased mRNA abundance. Other mutations in these domains showed increased mRNA abundance, suggesting complex regulation of *CBS* mRNA expression linked to other nucleotide or codon features.

### Differential mRNA abundance is consistent with codon-mediated stability and identifies variant effects undetected at the protein level

In yeast, zebrafish, *Xenopus*, and human cells, mRNA decay and translation efficiency are partially explained by codon optimality [47–52], where optimal codons are defined as those enriched in transcripts with longer mRNA half-lives and/or increased translation efficiency. While previous studies predominantly relied on reporter assays to assess the impact of codon optimality on these gene expression phenotypes, our approach enabled testing the relationship in the context of a native transcript coding sequence. To test if any *CBS* variants alter mRNA abundance through codon-mediated mRNA stability, we compared the difference in tRNA abundance and the codon stability coefficient (CSC) [50, 53] between reference and alternate codons to the magnitude of differential mRNA abundance (**Methods**).

tRNA abundance exhibited a modest correlation with the mRNA abundance fold change for variants down in mRNA (0.49 and 0.31 Spearman correlation for hydro-tRNAseq [54] and mim-tRNAseq [53], respectively) (**Figure 5B**, **Supplementary Figure 6C**). Further, the difference in CSC [50] between alternate and reference codons was lower for variants with decreased mRNA abundance compared to variants with increased abundance, indicating changes to less-stable codons may reduce mRNA levels (**Figure 5C**, one-sided Wilcoxon rank sum test p=0.029). Aside from the mutations at C52 and C272, important structural residues of CBS [55], mutations to cysteine (A38C, T135C, P138C) and from cysteine (C109G/T) exhibited effects consistent with its low codon stability [50] (**Supplementary Table 1**). By comparing each differential variant to a non-significant variant leading to the same amino acid change, we found nine variants had a CSC difference in the expected direction (**Figure 5D**) compared to four variants in the opposite direction (binomial test for 9 successes of 13, p=0.046). Notably, changes to the valine codon UUG, phenylalanine codon UUU, and arginine codon AGG may be candidates for codon-mediated reduction in mRNA abundance (**Supplementary Table 1**). Our results suggest codon optimality partially explains mRNA abundance effects for at least some missense variants.

Finally, we integrated paired yeast functional complementation data for the *CBS* variant library [10] and found 61/108 (56.5%) of variants called by both methods (FDR < 0.15) showed a consistent directional effect (e.g. low mRNA, low fitness) at a fitness score cutoff of 0.7 (score range 0-1, binomial test p=0.074). 16.9% of missense variants (14/83) with low mRNA abundance were at amino acid positions implicated in CBS deficiency. Of variants with decreased mRNA abundance, low fitness was confirmed for H65R/V, K119F, I122F, T135C, I142W, I143R, E144L, while other variants showed a more modest reduction of fitness (A38C, K39V, P49K, R58I, H66F, H67T, E128I, P138C) (**Supplementary Table 1**). Our results highlight the utility of mRNA abundance readouts to complement protein abundance and activity data. Together, we find variant effects on mRNA abundance are partially explained by codon-mediated stability and may diverge from yeast functional complementation readouts.

## Discussion

The explosion of saturation mutagenesis studies [2,5–7,9,10,56–58], facilitated by next generation tools to measure molecular phenotypes [4,5,29,59], prompts a need for an analysis solution that is extensible to multiple experimental designs. We created satmut_utils to fill this gap by providing simulation and variant calling in multiple amplicons simultaneously for both amplicon and RACE-like methods.

With satmut_utils ‘sim’, we conducted the first benchmarking analysis of MAVE variant callers and trained error correction models to achieve high variant calling performance. We further implemented several read preprocessing strategies (primer masking, consensus deduplication), which act synergistically with error correction models to improve specificity. Our goal with satmut_utils ‘call’ was to enable primary variant calling analysis to accurately resolve low-frequency SNPs and MNPs. The previously developed software methods for analysis of MAVEs [14–16] do not easily scale for large genes and are generally tailored to pre-/post-selection designs. Thus, we developed a general solution that makes limited assumptions about experimental design, and focuses on accurately identifying and quantifying variants prior to statistical inference.

One limitation of satmut_utils is that it is not compatible with barcode-sequencing, wherein a barcode (i.e. randomer) is separated from the mutagenesis region and linked to a specific variant. While this method simplifies variant calling and quantification, it requires initial sequencing for barcode assignment, which increases cost and time. Further, compared to direct variant calling, barcode-sequencing may lead to regulatory changes due to molecular linking between the barcode and the gene of interest.

We demonstrate the utility of our software solution by mapping coding variant effects on mRNA abundance for the *CBS* gene. Building on prior clinical and functional data [10], we assayed variant effects on *CBS* mRNA abundance in human cells and found several variants at important CBS structural residues with low mRNA abundance. Our results are consistent with a recent study that employed saturation genome editing of *BRCA1* to uncover hundreds of SNPs with differential mRNA abundance, and supported the notion that variants at key structural residues can lead to low mRNA abundance [60].

Both synonymous and nonsynonymous variants may have strong effects on expression through translation regulation, codon optimality, and alteration of mRNA secondary structure [61–67]. Optimal codons tend to be enriched in regions encoding buried, non-solvent accessible residues [68], which may explain our observation of a dependence of the magnitude of variant effect on position in the coding sequence. We also identified seven variants that converted to a non-optimal cysteine codon (UGU) and exhibited low mRNA abundance, consistent with its low codon stability [50]. We speculate mutations causing CBS deficiency may negatively feedback on *CBS* mRNA expression via reduced biosynthesis of cysteine to compound the deleterious effect of low stability of the UGU codon. Further work is needed to quantify the extent to which codon optimality modulates expression of endogenous transcripts [60] as opposed to reporter constructs, and satmut_utils is poised to support such studies.

Here we analyzed coding variant effects on mRNA abundance, but analysis of other MAVE data is possible. For example, yeast functional complementation assays [4, 10] and FACS-based assays to measure protein abundance [5,7,9] are plausible applications for satmut_utils analysis.

## Conclusions

We offer satmut_utils as a flexible solution for variant simulation and variant calling in saturation mutagenesis experiments. The satmut_utils package is unit-tested, well-documented, and available on GitHub and PyPi. Our method supports two different library preparation methods and incorporates state-of-the-art error correction through read pre-processing and machine learning models. Further, satmut_utils uses standardized input and output files and is compatible with existing statistical inference tools. In conclusion, satmut_utils is a complete solution for analysis of multiplexed assays of variant effect, and will motivate novel assays based on targeted DNA- and RNA-sequencing.

## Methods

### Code availability

Source code, installation instructions, and documentation is available on GitHub: https://github.com/CenikLab/satmut_utils. A Python package is available on PyPi: https://pypi.org/project/satmut-utils/. See the satmut_utils manual for details on usage and algorithmic design of the ‘sim’ and ‘call’ workflows.

### Secondary analysis

Secondary analysis of satmut_utils results, including benchmarking and error correction modeling, used a pre-release version of satmut_utils with accessory scripts, located at https://github.com/ijhoskins/satmut_utils. Supplementary Tables, Additional files, and reference files used in analysis are available at https://github.com/ijhoskins/satmut_utils_supplementary. All analyses were carried out on an AMD Ryzen Threadripper 3990X 64-Core Processor with Ubuntu OS.

### satmut_utils ‘sim’ workflow

The satmut_utils ‘sim’ workflow takes a Variant Call Format (VCF) and alignment (BAM) file with paired reads as input, and generates variants in the reads at specified frequencies. Outputs are a VCF of true positive (truth) variants and counts, along with edited reads (FASTQ). ‘sim’ is comprised of three overall steps: 1) a single samtools ‘mpileup’ call [69, 70] is made to query reads at each position in the target region. The number of fragments to edit and the read positions to edit are determined for each variant based on specified frequencies in the input VCF. ‘sim’ employs a heuristic to sample reads for editing at each target position while prohibiting variant conversion (the merging of edited variants with nearby errors). 2) With these edit configurations, variants are edited into read pairs and written as raw reads in FASTQ format by ‘samtools fastq’. 3) The raw reads are re-aligned with bowtie2 [71] *global* alignment mode to generate valid CIGAR and MD tags, which are required for visualization of edited reads in genome browsers.

### satmut_utils ‘call’ workflow

satmut_utils ‘call’ utilizes cutadapt [72] for adapter and 3’ base quality trimming, followed by an optimized, paired-end *local* alignment to the transcript reference using bowtie2 [71] with the following parameters: ‘--maxins=1000 --no-discordant --fr --mp 4 --rdg 6,4 --rfg 6,4’. If consensus deduplication is requested, this step directly follows alignment. Then, if a primer BED file is provided, primer base quality masking is performed. Following read preprocessing, filtering on base quality, read edit distance, and min supporting counts is applied during variant calling. Variant calls are made by iterating over filtered read pairs, finding mismatches with mate concordance, extracting quality features, and *writing results for each mismatch participating in a primary variant call*. (This should be considered when counting records from output files). Fragment coverage depth is reported in bedgraph format. To validate satmut_utils ‘call’, we generated *in silico*, error-free, paired-end RNA reads and then introduced 187 SNPs and MNPs, each at 10 read pairs in 10,000 (0.1%). Tuning of bowtie2 InDel penalties was required to achieve 100% recall for MNPs (Supplementary Figure 1E).

### satmut_utils primer base quality masking

If a primer BED file is provided, alignments are intersected with primers with ‘bedtools intersect -bed -wa -wb’. The resulting BED file is processed with ‘bedtools groupby -o collapse’ to group the intersecting primers for each read, and primers which originate the read are determined by the following criteria: 1) the read 5’ end begins within the aligned coordinates of the primer, or starts within a buffer upstream of the primer 5’ end (relative to strand); 2) the read 3’ end stops within the aligned coordinates of a primer on the opposite strand, or stops within a buffer upstream of the primer 5’ end (relative to strand). The buffer is 15 nt for amplicon methods and 3 nt for RACE-like methods. (These rules ensure masking for 3’ base-quality trimmed reads and reads with slight differences in alignment start and stop coordinates, for example due to incomplete primer synthesis or alignment clipping). Subsequences for originating primers are masked in the reads by setting the base qualities of these read segments to 0. These bases are not subsequently considered for variant calling and fragment coverage enumeration.

### satmut_utils UMI-based consensus deduplication

Unique molecular indexes (UMIs) are extracted to the read header prior to adapter and 3’ base quality trimming. Following alignment, reads are grouped by [UMI x R1 POS] with UMI-tools [73] default directional adjacency method and the --paired, --ignore-tlen flags. Group ID tags are copied from R1 alignments to R2 alignments, paired reads are combined and sorted by read name, and then R1 and R2 are consensus deduplicated using a majority vote at each aligned position in the UMI group. In base call ties (two duplicates), if one base call matches the reference base, the reference base is used for the consensus. Otherwise, the base call with the higher base quality is used, thereafter defaulting to random choice.

### Data preprocessing for benchmarking

Reads originating from a single *CBS* negative control amplicon (tile 6) from the wild-type, non-selected condition [10] were selected for simulation. To meet dms_tools2 and Enrich2 input requirements, reads were preprocessed using an accessory script (https://github.com/ijhoskins/satmut_utils/blob/satmut_utils_dev/src/scripts/run_design_conversion.py). Preprocessing comprised several steps: 1) reads were locally aligned; 2) any hard- or soft-clipped reads, unpaired singletons, and reads with InDels were filtered out; 3) reads were modified to start and end flush with codons by trimming and/or appending reference sequence; 4) for dms_tools2, 12 nt unique molecular indices (UMIs) were added to the 5’ end of both R1 and R2, enforcing unique UMIs for each read pair.

An accessory script (https://github.com/ijhoskins/satmut_utils/blob/satmut_utils_dev/src/scripts/run_ec_data_generator.py) was used to generate the benchmarking dataset. 281 SNPs and MNPs were simulated in the preprocessed negative control (NC) alignments for tile 6 of *CBS*, using frequency parameters estimated from satmut_utils variant calls (-m 2 -q 30 -e 10-s NNK) across all tiles of the mutagenized, non-selected condition [10]. In addition, the proportion of SNPs in the truth set was set at 0.25. To balance the number of true and false positive labels, the number of variants to edit was determined by a heuristic that samples variants until the number of component mismatches comprising these variants equals the number of false positive mismatches in the NC library. The number of false positive mismatches in the NC library was determined by satmut_utils call using the following parameters: ‘-m 1 -q 30 -e 10 -s NNK’.

Configurations and quality parameters for benchmarking were as follows. CBS_TILE6_SEQ = “GACGTGCTGCGGGCACTGGGGGCTGAGATTGTGAGGACGCCCACCAATGCCAGGTTCGACTCCCCGGAGTCACACGTGGGGGTGGCCTGGCGGCTGAAGAACGAAATCCCCAATTCTCACATCCTAGACCAGTACCGCAACGCCAGCAACCCC”

1. DiMSum: ‘--stranded=T -q 30 -m 10 -u coding --mutagenesisType=codon --indels=none --mixedSubstitutions=T -s 1 -t 4 -w $CBS_TILE6_SEQ’
2. dms_tools2: ‘--alignspecs 19,132,31,34 --bclen 12 --bclen2 12 --chartype codon --maxmuts 10 --minq 30 --minreads 1’
3. Enrich2: {"filters": {"avg quality": 20, "max N": 10}; "variants": {"max mutations": 3, "min count": 2, "use aligner": false, "wild type": {"coding": true, "reference offset": 534, "sequence": $CBS_TILE6_SEQ}}} The “Basic” mode was used (variant calling on R1 only), and variants to the unknown base N were filtered out.
4. satmut_utils: ‘-m 2 -q 30 -e 10 -s NNK’

A *post-hoc* filter was applied to select variants with MATCHES_MUT_SIG=True.

### Generation of error correction validation datasets

The same accessory script used for generation of the benchmarking dataset was used for generation of four ‘sim’ datasets. To estimate error correction parameters, we ran satmut_utils ‘call’ on each NC library with the parameters ‘-m 1 -q 30 -e 10 -s NNK’ to count false positive mismatches in the control alignments. satmut_utils ‘call’ was also ran on an input source-matched, mutagenized library with the same parameters, except with a min count of 2 (-m 2). NC and mutagenized satmut_utils summary.txt files, along with the trimmed NC alignments, were used as inputs to the script. To complete each simulated dataset, satmut_utils ‘call’ was ran on the output FASTQs, with the same parameters as NC libraries. Each simulated dataset (N=4) comprised thousands of true positives (min 4850, max 7859) and thousands of false positives (min 4682, max 6463).

### Data postprocessing and error modeling Custom R functions in

(https://github.com/ijhoskins/satmut_utils/tree/satmut_utils_dev/src/prototype) were used to postprocess and model the resulting simulated datasets. The packages data.table [74], ggplot2 [75], cowplot [76], viridis [77], and ggsci [78] were used for data processing and graphics. The following packages were used for modeling: leaps [79], caret [80], e1071 [81], class [82, 83], randomForest [84], gbm [85], and glmnet [86].

Variant calls in each simulated dataset within the mutagenized target region, and with frequency < 0.3, were selected for modeling. Five classifiers were trained: gradient boosted machine (decision trees); generalized linear model (binomial family) with elasticnet regularization; k-nearest neighbors; random forest; and support vector classifier. Performance was evaluated by nested 10-fold cross-validation (CV), selecting 20% of each fold’s training data for hyperparameter tuning with caret::train. Additionally, for k-nearest neighbors, the number of features was tuned in each fold with best subset selection (leaps::regsubsets) by 5-fold CV, using between 3 and 10 features. For predictions of all models, a probability cutoff of 0.5 was used. The feature importance metric (mean decrease in accuracy) was determined by passing importance=TRUE during random forest training and subsequently calling randomForest::varImpPlot(type=1).

### Differential abundance analysis

Variant calls were filtered by several sequential steps prior to differential abundance analysis. First, variant frequencies were adjusted by subtracting the log_10_ variant frequency in the NC library from the frequency of corresponding variants in the mutagenized libraries. Then, candidate variants were selected in sequential order by the following criteria: 1) variant is within the mutagenized target region; 2) variant matches the NNK codon mutagenesis signature; 3) variant is a single-codon change; 4) SNP variant count >= 2 and MNP variant count >= 1; 5) no strong strand bias (RACE-like method only, nucleic acid strand count difference <= 64); 6) no variants with false positive RF predictions in all replicates (probability cutoff 0.49); 7) variant is observed in all replicates. Additionally, for amplicon data, one sequencing library with possible bottlenecking (gDNA replicate 3, Supplementary Figure 4A) was dropped and replaced with the plasmid library sample to achieve three replicates. Amplicon method gDNA replicate 3 was warranted for exclusion as it formed its own cluster from other gDNA and cDNA replicates by hierarchical clustering analysis.

For filter 6, model training datasets were generated as described, and a RF model was trained on the following features: log_10_ frequency, variant type (SNP, di-nt MNP, tri-nt MNP), matches mutagenesis signature, substitution (e.g. A>G,T>C; six factor levels), upstream reference nt, downstream reference nt, R1 and R2 median supporting base qualities, R2 median supporting read position, R2 median supporting edit distance. For the RACE-like model, R1 and R2-specific features (base quality, read position, read edit distance) were additionally provided for each sample strand (R1+, R1-, R2+, R2-), along with the sample strand count difference. For read position and edit distance features, only R2 was used due to collinearity with the corresponding R1 features. RACE-like model training also required na.action=”na.roughfix” to handle NAs in training data due to count observations on only one sample strand.

To determine variants with differential abundance between mRNA and gDNA, we used limma-trend [87] with empirical Bayes moderation and Benjamini-Hochberg multiple test correction. gDNA readout serves to normalize for library abundance and recombination efficiency of each variant. Relative changes in total RNA thus reflect variant effects on population-wide, steady state mRNA abundance.

### Replicate analysis

For Figure 4E-G, variant calls were processed as described for differential abundance analysis with the exception of the last criterion (#6, variant observed in all replicates). Instead, variants observed in only one replicate were filtered out. For Figure 4G, due to lower depth of coverage in RACE-like sequencing libraries, the median variant frequency of cDNA replicates was plotted for variants with a log_10_ frequency > -5.2 in amplicon gDNA libraries. This is the approximate limit of detection for the RACE-like method given the attained coverage.

### Comparison of library preparation methods

The theoretical number of possible calls in CBS tiles 2 and 4 (2676) was calculated empirically by counting all single SNPs, di-nt MNPs, and tri-nt MNPs that match a NNK mutagenesis signature for each codon in the *CBS* coding sequence. Unless otherwise noted, variant counts, frequencies, and cDNA-gDNA frequency difference (effect estimates) follow filtering as described in *Differential abundance analysis* and *Replicate analysis* and use the median for replicate summarization.

For variant effect comparison (Supplementary Figure 5C), variants determined significant in the amplicon method at FDR < 0.1 were assessed in RACE-like data. Many variants were at or below the limit of detection, so NAs were replaced with the approximate limit of detection (the maximum of the minimum variant frequency across all gDNA and cDNA replicates, log_10_ frequency -5.38).

### Analysis of tRNA abundance

Variants identified in the amplicon method at a FDR < 0.15 were used for analysis of tRNA abundance data to achieve better power for analysis. For mim-tRNAseq [53], the mean of counts was taken of the HEK293T duplicates. For hydro-tRNAseq [54], HEK293 counts were used directly. For both datasets, anticodon abundance for codons with Crick wobble base pairing (A-G and C-T) were added to the dataset. Then, the sum of isodecoder counts was taken for each codon and log2 transformed. The difference between the log2 sum of counts was calculated between the alternate (variant) and reference codons and compared to the log fold changes determined by limma.

### Analysis of codon stability coefficient (CSC) data

The same set of variants used in analysis of tRNA abundance data was used to test for differences in CSC for the ORFome in HEK293T cells [50]. For Figure 5C and D the difference between the alternate codon (Alt.) and reference codon (Ref.) CSC was computed. For Figure 5D, all pairwise distances of this difference between significant and non-significant variants (691 total variants, “delta delta”) was determined to define the null distribution. Variants with a distance in the lower quartile of this null distribution were filtered out, and variants in the upper quartile of the distribution were marked with an asterisk.

### Cell culture

HEK293T LLP iCasp9 Blast cells [29] were confirmed to be free of Mycoplasma and were cultured in Dulbecco’s Modified Eagle Medium (Thermo Fisher Scientific, 11995065) with 10% fetal bovine serum (Gibco, and 1% penicillin-streptomycin (Gibco, 15140122). Prior to recombination at passage 16, cells were selected for one week with 2 µg/mL Doxycycline (Sigma-Aldrich, D3072) and 10 µM Blasticidin (Gibco, A1113903) to enrich for cells with the integrated landing pad.

### *CBS* library cloning

A *CBS* variant library was generated as previously described [10]. The *CBS* entry library was transferred into pDEST_HC_rec_bxb_v2, a vector containing recombination sites for the HEK293T LLP iCasp9 Blast landing pad line, by a Gateway LR II reaction (Thermo Fisher Scientific, 11-791-020) following manufacturer’s recommendations. 1.5 µL of LR reaction was transformed into 25 µL Endura Electrocompetent cells (Lucigen, 60242), plated on Nunc Square Bioassay dishes, scraped in 6 mL LB Miller broth, and 3 mL resuspension was processed with the ZymoPURE II Plasmid Maxiprep Kit (Zymo Research, D4203). Library size was estimated at ∼540,000 species, or ∼30-fold coverage of each possible SNP or MNP variant in the *CBS* coding sequence.

### Stable expression of *CBS* variant library

20 µg of the *CBS* variant library (in pDEST_HC_Rec_Bxb_V2), along with an equal mass of Bxb1 recombinase (pCAG-NLS-HA-Bxb1) was transfected into three 15 cm dishes of HEK293T LLP iCasp9 Blast cells (passage 18, 65% confluency) using Lipofectamine 3000 (Thermo Fisher Scientific, L3000008), with volumes scaled based on 3.75 µL reagent per 6-well. 24 h later, cells were split 1:2 into 15 cm dishes. 48 h after transfection, at near full confluency, 2 µg/mL Doxycycline and 10 nM AP1903 (MedChemExpress, HY-16046), both solubilized in DMSO, were added for negative selection of non-recombined cells. The next day, dead cells were removed and recombined cells were grown out for an additional two days with fresh media containing Doxycycline and AP1903. Cells were recovered for one day by growth in media without Doxycycline and AP1903. Transcription was induced with 2 µg/mL Doxycycline for 24 h, cells were stimulated with fresh media for 3 h, and then harvested at 95% confluency.

### gDNA extraction

gDNA was extracted from approximately 3-4 million cells with the Cell and Tissue DNA Isolation Kit (Norgen Biotek Corp, 24700), including RNaseA treatment and eluting in 200 µL warm elution buffer.

### RNA extraction and cDNA synthesis

Approximately 3-4 million cells were solubilized with 1 mL QIAzol (Qiagen, 79306) and 0.2 mL chloroform in 5PRIME Phase-Lock Gel heavy tubes (QuantaBio, 2302830), according to the manufacturer’s recommendations. RNA was precipitated at -20 °C following the addition of 2 µL GlycoBlue (Thermo Fisher, AM9515) and 2.5 volumes of cold absolute ethanol. RNA was washed once with cold 70% ethanol then resuspended in 30 µL water. 10 µg total RNA was treated with DNaseI (NEB, M0303), then re-purified by the RNA Clean and Concentrator Kit (Zymo Research, R1015) and eluted in 15 µL water. RNA quality was assessed with the Bioanalyzer Eukaryotic RNA Pico kit (Agilent, 5067-1513). For each of six reactions, 2.5 µg DNaseI-treated total RNA was denatured at 65 °C for 5 min followed by RT primer annealing at 4 °C for 2 min, using 2 pmol pDEST_HC_Rec_Bxb_v2_R primer specific for the landing pad. See Supplementary Table 2.

Primed total RNA was included in six 20 µL SuperScript IV cDNA synthesis reactions (Thermo Fisher, 18090010) with SUPERase-In RNase inhibitor (Thermo Fisher, AM2696), and first-strand cDNA was synthesized by incubating at 55 °C for 1 h, followed by RT inactivation at 80 °C for 10 min. RNA was digested with addition of 5 U RNaseH (NEB, M0297) to the first strand cDNA synthesis reaction and incubation at 37°C for 20 min. One reaction was saved for amplicon library preparation, while the other five were saved for RACE-like (Anchored Multiplex PCR) library preparation.

### Amplicon library preparation

See Supplementary Table 2 for primers used in PCR1 and PCR2 of amplicon method library preparation, outlined below.

#### Landing pad amplification (PCR1)

2.5 µg of gDNA was amplified with Q5 polymerase (NEB, M0491) for 14 cycles in each of six 50 µL PCR reactions with 500 nM landing-pad-specific primers (pDEST_HC_Rec_Bxb_V2_F, pDEST_HC_Rec_Bxb_v2_R) flanking the entire *CBS* insert (∼1.7 kb), and including the high GC enhancer reagent. The cycling parameters were: initial denaturation at 98 °C for 30 s; 3-step cycling with denaturation at 98 °C for 10 s, anneal at 65 °C for 30 s, extension at 72 °C for 1 min; final extension at 72 °C for 2 min.

Products were pooled, resolved on a 0.8% agarose/TAE gel, visualized with 1x SYBR Gold, and extracted from the gel using the Macherey-Nagel Nucleospin Gel and PCR Cleanup Kit (Takara, 740609) with 15-25 µL 70 °C elution buffer.

#### Coding sequence amplification (PCR2)

500 pg of the gDNA and cDNA PCR1 products (landing pad insert) was amplified for each of two CBS tiles (CBS_2_v2, CBS_4_v2 primer pairs) in a 50 µL NEB Q5 reaction (NEB, M0491) with high GC enhancer for 8 cycles, following the same cycling parameters as for PCR1.

#### Illumina adapter addition (PCR3)

Products for tile 2 and 4 amplicons were cleaned up with the Nucleospin Gel and PCR Cleanup Kit (Takara, 740609), eluted in 30 µL 70 °C buffer, and then mixed together at equal volumes (5 µL each) and input into a final NEB Q5 reaction for 8 cycles with the same formulation as PCR2 but using NEBNext Multiplex Oligos for Illumina Dual Index Primers Set 1 (NEB, E7600S) according to manufacturer’s recommendations (65 °C annealing). Final library was purified with the Nucleospin Gel and PCR Cleanup kit and eluted in 30 µL 70 °C buffer.

#### RACE-like library preparation

Anchored Multiplex PCR libraries were generated with modifications following the initial strategy [20]. Briefly, for cDNA libraries, double-strand cDNA was first synthesized, and gDNA and double-strand cDNA inputs were sheared prior to library preparation. Libraries were prepared using the ArcherDX, Inc. (now Invitae) LiquidPlex library preparation kit with a custom *CBS* primer assay (https://archerdx.com/research-products/custom-panels/). See Supplementary Tables 2 and 3 for primer sequences.

#### Second strand cDNA synthesis

Second strand cDNA synthesis was carried out with 1^st^ strand cDNA from each of five reactions converting 2.5 µg total RNA and digesting with 5U RNaseH. 2 pmol landing-pad-specific forward primer (pDEST_HC_Rec_Bxb_v2_F) and 1 µL Q5 polymerase (NEB, M0491) were added to 20 µL of each 1st strand cDNA reaction and incubated as follows: 95 °C denaturation for 30 s, followed by three cycles of 1) 55 °C anneal for 30 s and, 2) 72 °C extension for 3 min, for a total of three linear primer extension cycles. Reactions were pooled and cleaned up with the Nucleospin Gel and PCR Cleanup kit with 1:1 buffer NTI dilution and elution with 15 µL 70 °C elution buffer.

#### DNA fragmentation

gDNA was extracted from 5 million cells with the Cell and Tissue DNA Isolation Kit (Norgen Biotek Corp, 24700), including RNaseA treatment and eluting in 200 µL warm elution buffer. gDNA and double-strand cDNA were fragmented using the Covaris S2 with the following parameters: microTUBE AFA Fiber Snap-Cap, 10% duty cycle, 5 intensity, 200 cycles per burst, 1 min treatment. Sheared DNA inputs were brought up to 50 µL with ultrapure water.

#### PCR enrichment and library preparation

Libraries were prepared from 1-1.5 µg sheared gDNA or double-stranded cDNA according to the manufacturer recommendations (ArcherDX, PRO027.4), using 15 cycles for both PCR1 and PCR2 with custom *CBS* primers (Supplementary Table 3).

### Quantification and size-selection of final libraries

Final libraries were quantified using the KAPA Library Quantification Kit for Illumina (Roche, KK4873) according to the manufacturer’s recommendation, using 1:10,000 dilution of libraries and size-correction using an average fragment size of 300 nt. qPCR was performed on the Applied Biosystems ViiA 7 instrument. The final pool was size-selected using Sage Bioscience BluePippin 2% gel (marker V1 and/or V2) to select fragments from 200-400 bp. Following size-selection, the final pool was quantified by both the High Sensitivity DNA Kit (Agilent, 5067-4626) and Qubit High Sensitivity DNA assay (Thermo Fisher, Q32851).

### Preparation of negative control libraries

Negative control libraries and mutagenized libraries used for estimating true variant frequencies were prepared from various sources (HEK293T total RNA, plasmid DNA) with several methods. See protocols for each library under the GEO submission. Primer sequences for all CBS amplicons are provided in Supplementary Table 4.

### Next-Generation Sequencing

Libraries were sequenced 2 x 150 (paired-end) at MedGenome, Inc. on the Illumina HiSeq X platform. RACE-like (Anchored Multiplex PCR) libraries were re-sequenced on the Illumina NovaSeq 6000 and FASTQs were concatenated prior to analysis. 5-10% PhiX was included during sequencing.

## Declarations

### Ethics approval and consent to participate

Not applicable.

## Consent for publication

Not applicable.

## Availability of data and materials

satmut_utils is available on GitHub at https://github.com/CenikLab/satmut_utils. See the package setup.cfg file for software dependencies and versions. Accessory scripts used for secondary analysis, benchmarking, and error correction modeling are available at https://github.com/ijhoskins/satmut_utils/tree/satmut_utils_dev/src/scripts.

Raw sequencing data is available at GEO under accession GSE201057. Simulated datasets are available from the corresponding author on request.

## Authors’ contributions

**IH**- Conceptualization, Methodology, Software, Validation, Formal analysis, Investigation, Visualization, Writing- Original Draft. **SS**- Investigation. **AC**- Investigation. **CC**- Conceptualization, Writing- Review and Editing, Supervision, Project administration, Funding acquisition. **FR**: Resources, Supervision.

## Acknowledgements

We would like to thank Doug Fowler lab for kindly providing the HEK293T LLP iCasp9 Blast cell line. We thank Shilpa Rao and Sun-Young Kim for help with generating negative control sequencing libraries. We also thank for generating a primary human transcriptome for use with satmut_utils.

---

**Supplementary Figure 1:**
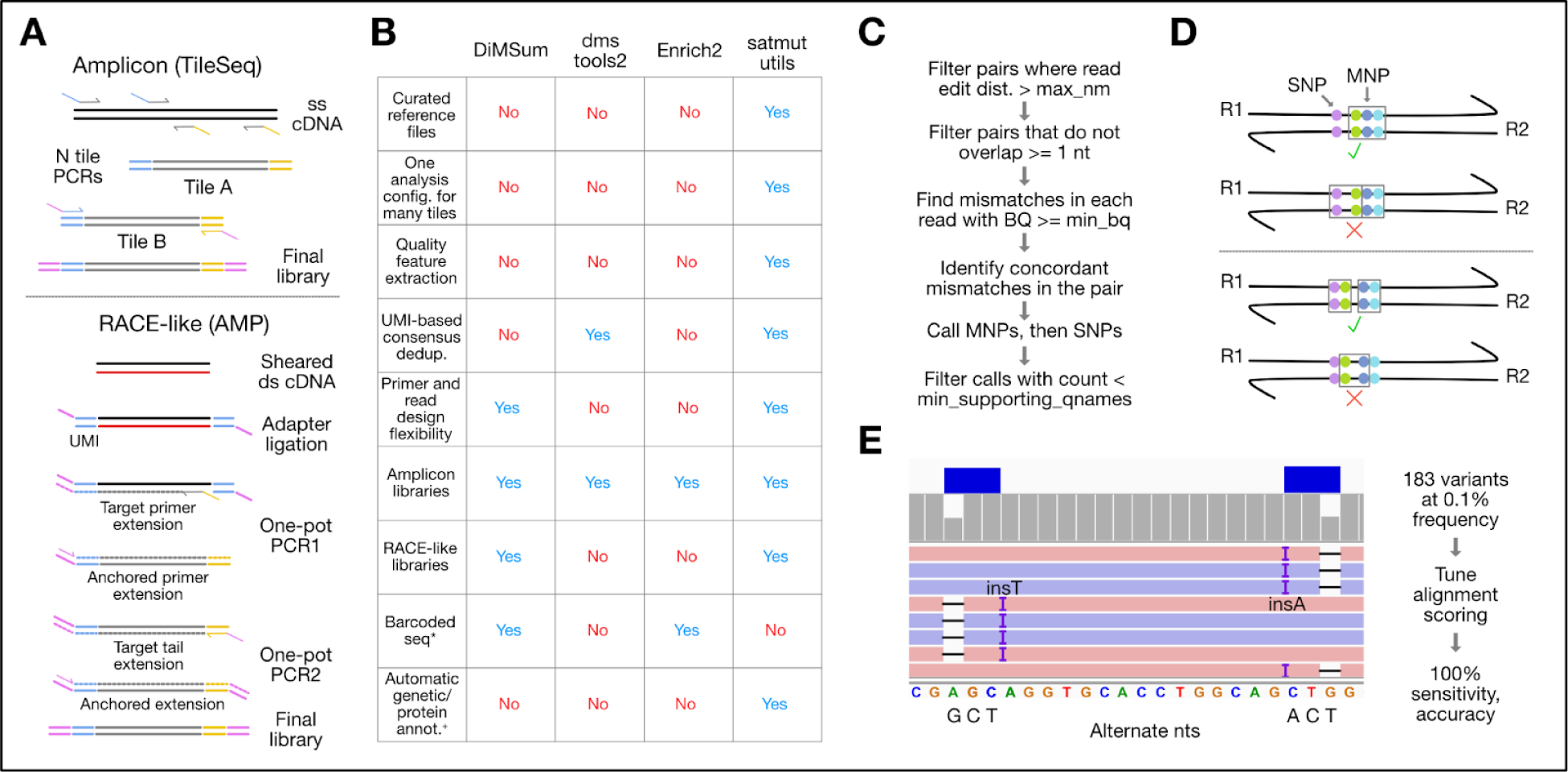
**A) Comparison of amplicon and RACE-like library preparation methods.** Pink segments indicate the first half of the adapters (Illumina P5/P7 sites). Blue and yellow segments indicate the rest of the adapter, containing read primer sites and possibly a unique molecular index (UMI). TileSeq is an amplicon method [4]. Anchored Multiplex PCR (AMP) [20] is a RACE-like method. **B) Feature comparison of MAVE variant callers.** Blue text indicates software utility/flexibility while red text indicates unsupported features. *Barcoded-seq refers to estimating variant abundance through counting a linked unique barcode. ^+^Codon and protein change annotations are reported with correct coding sequence positions, requiring no manual configuration of offsets. **C) Variant calling algorithm.** Read pairs are filtered with max edit distance (dist.) and min base quality (BQ) parameters before finding concordant mismatches (same base call in R1 and R2). **D) Schematic of MNP calling algorithm.** Gray boxes denote the MNP span. In cases where there are multiple mismatches within the window (--max_mnp_window), satmut_utils prioritizes contiguous mismatch runs. In one hypothetical case (top), a mismatch precedes a contiguous run of three mismatches. The compact run is called as a tri-nt MNP, and the preceding mismatch is called as a SNP. In a second case (bottom), two compact runs- each with two mismatches- are spaced by one base matching the reference. Each run is called as a di-nt MNP. **E) Example of tri-nt MNPs aligning as InDels.** Under default bowtie2 alignment parameters (--rdg/--rfg 5,3), MNPs may be aligned as InDels. After adjustment of the scoring parameters (--rdg/--rfg 6,4), MNPs aligned as contiguous mismatches (Methods).

**Supplementary Figure 2:**
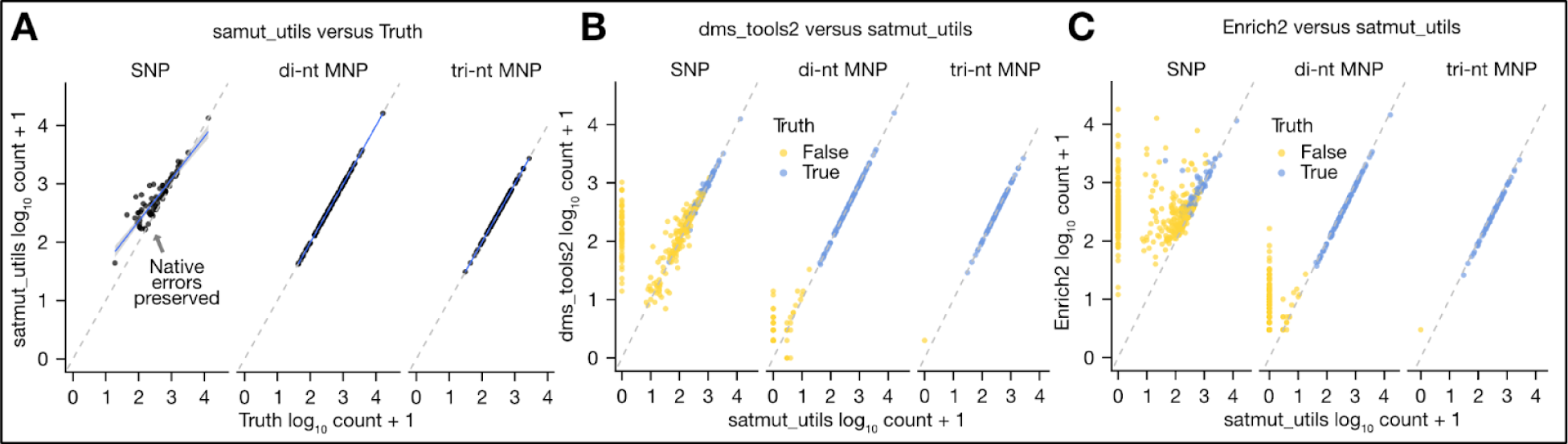
Comparison of variant callers for nucleotide changes. **A) satmut_utils count accuracy.** Simulated truth counts are compared to satmut_utils reported counts. Deviation for SNPs is due to preservation of native errors during simulation. Dotted gray lines indicate equivalence. Blue lines show the slope of a linear regression fit between truth and observed counts, with 95% confidence intervals in gray. **B) satmut_utils comparison to dms_tools2.** Counts for true (blue) and false positive (yellow) variants are shown. **C) satmut_utils comparison to Enrich2.** Counts are shown as in B. Higher false positive variants for Enrich2 is partly due to use of its Basic mode, which uses only R1 for variant calling. Enrich2 Overlap mode led to a high proportion of unresolved calls, which precluded analysis (see Additional File 1).

**Supplementary Figure 3:**
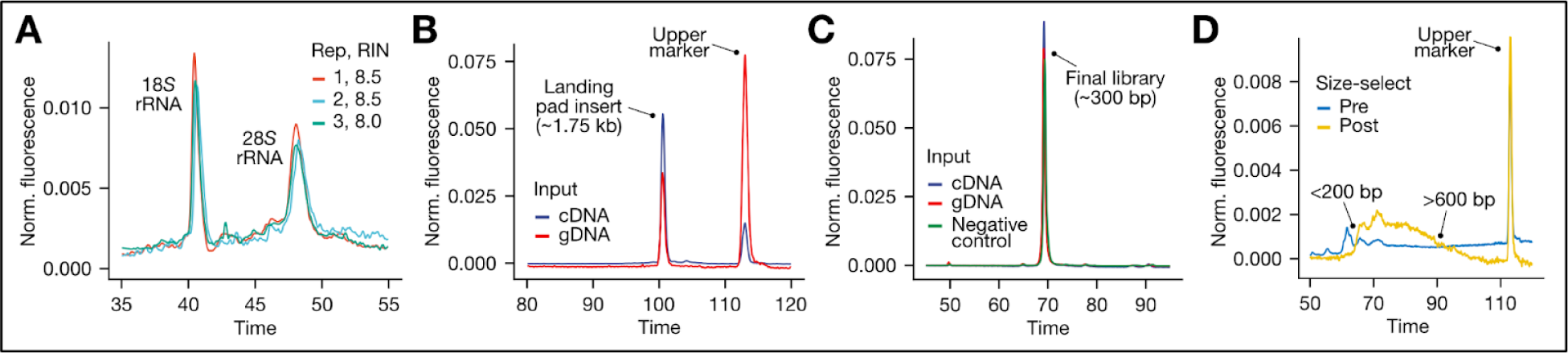
Library preparation quality control. For all panels, normalized (Norm.) fluorescence was computed by dividing by the sum of fluorescence across the displayed Time range. Panel A shows results from the Agilent Eukaryotic RNA Pico kit. Panels B-D show results from the Agilent High Sensitivity DNA kit. For panels B and C, traces indicate the mean of biological replicates. **A) Quality of biological replicate total RNA**. DNaseI-treated total RNA was assayed and the 18*S* and 28*S* rRNA peaks for each replicate (Rep), with RNA integrity number (RIN), are shown. **B) Confirmation of intermediate products for the amplicon method**. PCR1 was performed to enrich the landing pad insert (*CBS* coding sequence) from gDNA and cDNA prior to PCR2 for tiled amplicons. **C) Final library confirmation for the amplicon method**. Analysis of final libraries (PCR3) confirmed a specific product of the expected size (∼150 bp insert plus adapters). **D) Final library confirmation and size-selection for the RACE-like method**. Final gDNA and cDNA libraries were pooled and assayed before and after size-selection (Methods). Exclusion of incompletely-adapted library and short (<50 bp) or long (>450 bp) inserts is denoted.

**Supplementary Figure 4:**
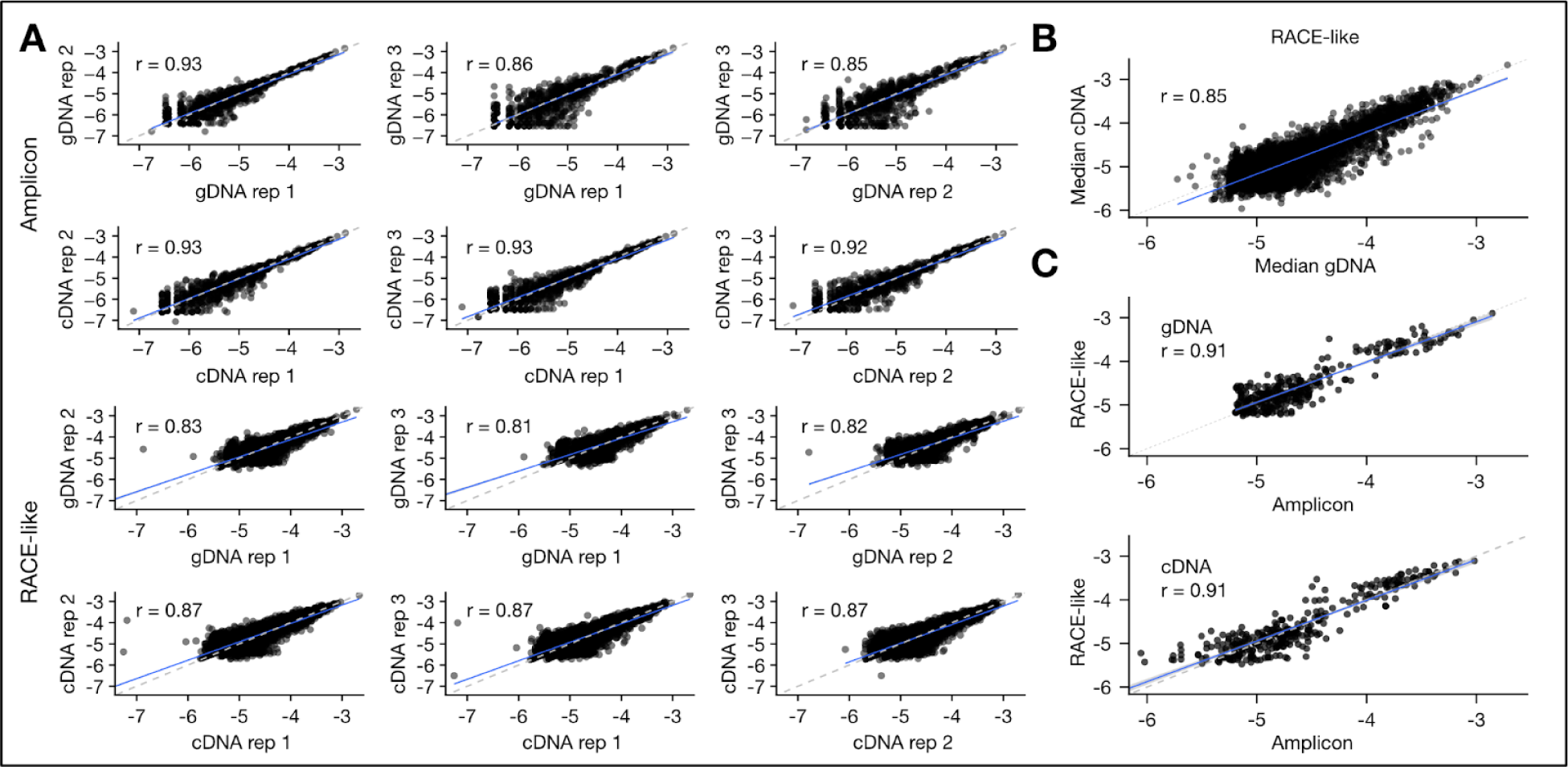
*CBS* variant frequency correlations for amplicon and RACE-like methods. In all panels, log_10_ variant frequencies are plotted after filtering out variants found in only one gDNA or cDNA library replicate. Pearson’s correlation coefficient (r) is indicated. Grey dotted lines indicate equivalence. Blue lines are a linear regression fit, with grey shading indicating 95% confidence intervals. **A) Biological replicate reproducibility**. gDNA and cDNA variant frequencies are shown for replicate cell lines independently recombined with the *CBS* variant library. **B) Correlation between gDNA and cDNA for RACE-like libraries**. The median frequency was computed among gDNA and cDNA replicates prior to comparison. **C) Variant frequency correlation between methods**. Due to lower depth of coverage for RACE-like libraries, variants with log_10_ gDNA frequencies greater than -5.2 were selected for comparison to the RACE-like method (Anchored Multiplex PCR) [20]. Replicate summarization used the median. Top panel compares gDNA libraries, and bottom panel compares cDNA libraries.

**Supplementary Figure 5:**
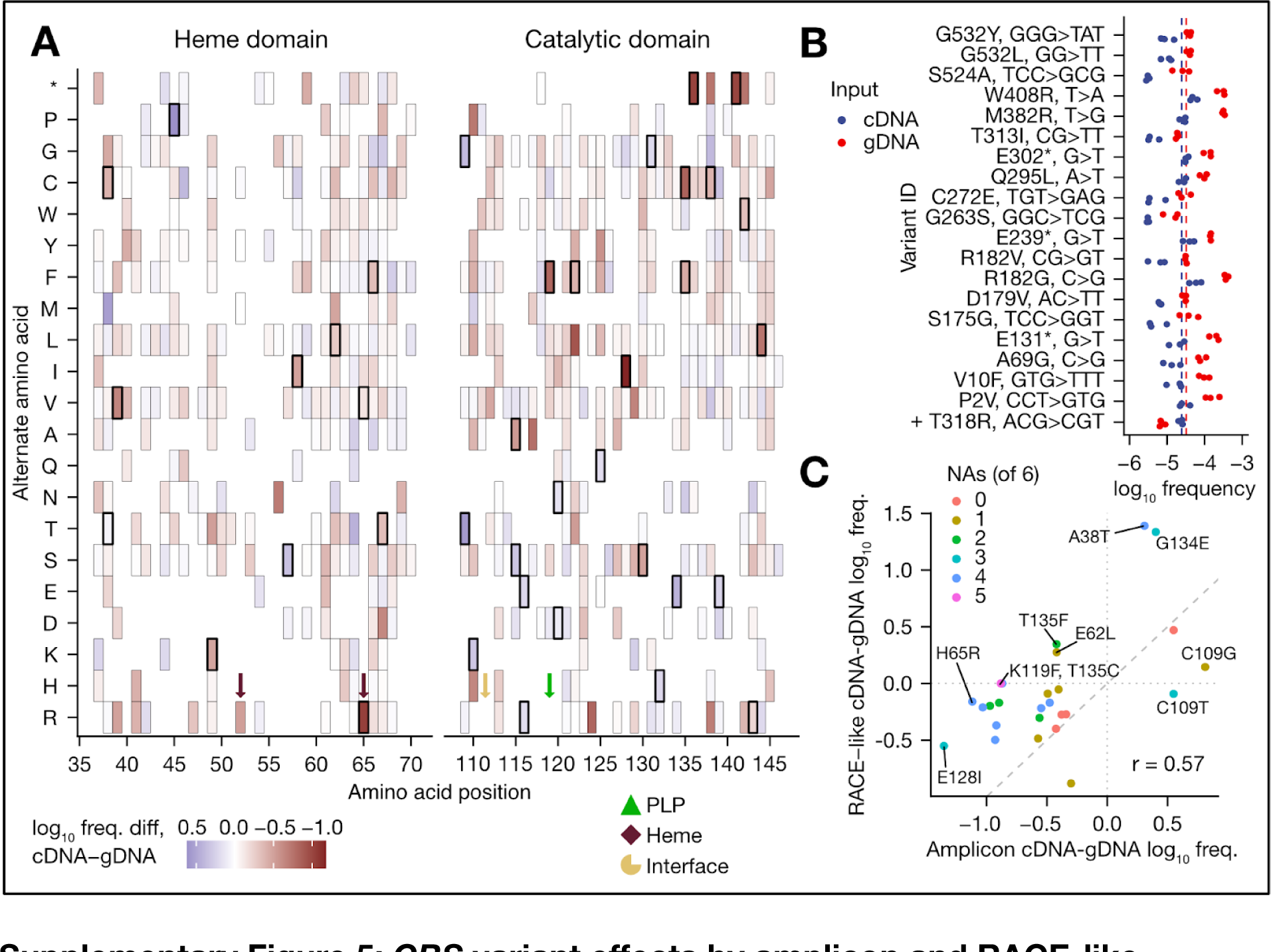
*CBS* variant effects by amplicon and RACE-like methods. **A) Amplicon method variant effect map**. log_10_ frequencies for variants leading to the same amino acid change were summarized; fill is the median log_10_ frequency difference between cDNA and gDNA. Bold borders indicate amino acid changes with at least one significant variant (codon change) with an effect in either direction. Arrows demarcate residues with heme binding (C52, H65), pyridoxal-5’-phosphate (PLP) binding (K119), or a location at the dimerization interface (111-112). **B) CBS variant differential abundance by RACE-like method.** y-axis labels denote the amino acid change and the nucleotide substitutions for variants with significant differential abundance at a FDR < 0.1). Red and blue dotted lines represent the median of each input source for all variants. (+) denotes one variant up in RNA. **C) Correlation of variant effects between amplicon and RACE-like methods.** Variants significant by the amplicon method (FDR < 0.1) are plotted if present in at least one replicate by the RACE-like method. The median log_10_ frequency difference between cDNA and gDNA is plotted following replacement of NAs with the approximate limit of detection (log_10_ frequency -5.38, Methods). Color indicates the number of RACE-like replicates in which the variant was observed. Pearson’s correlation coefficient is shown.

**Supplementary Figure 6:**
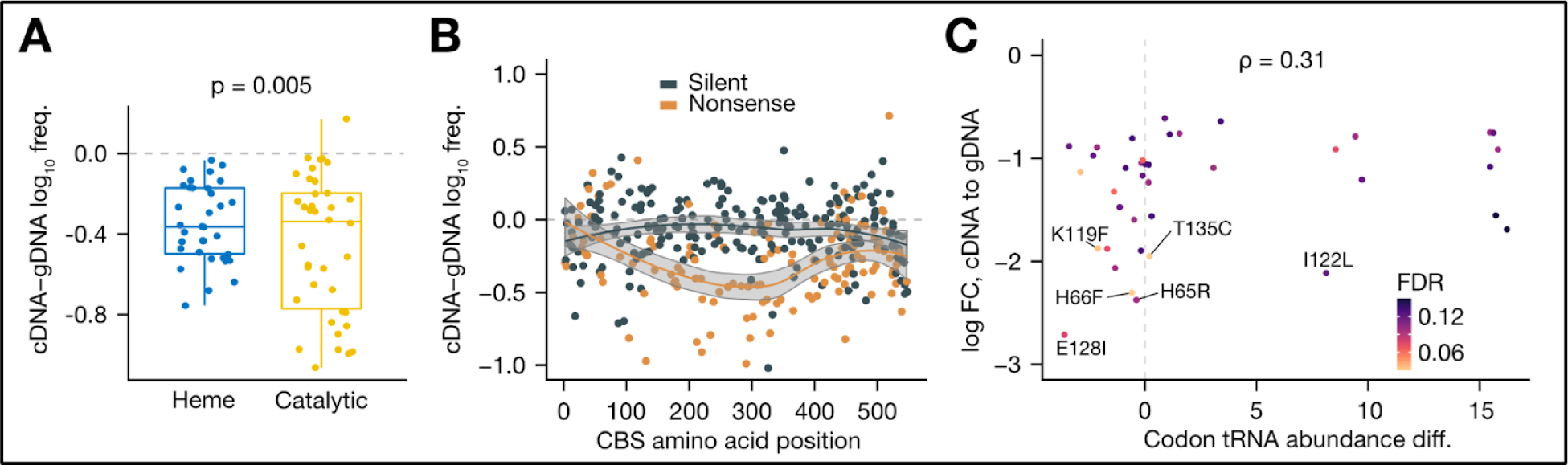
CBS variant effect dependence on position and comparison to tRNA abundance. In all panels, a grey dotted line indicates no change in variant effect or comparative metrics. **A) Higher variance of effects in the catalytic domain**. The minimum log_10_ difference between cDNA and gDNA for data from the amplicon method was computed at each targeted position and tested by a rank-based Brown-Forsythe Levene-type test. **B) Nonsense variant effects depend on location**. The difference in median log_10_ frequency between cDNA and gDNA for silent and nonsense variants in the RACE-like method is plotted across the *CBS* coding region. Lines represent local regression fit, with 95% confidence intervals in shaded grey. **C) tRNA abundance correlation with magnitude of variant effect**. The difference in tRNA abundance (Alt. codon - Ref. codon log2 sum of counts; HEK293T mim-tRNAseq [53]) is plotted against the log FC for variants identified by the amplicon method (FDR < 0.15). Spearman rank correlation is shown. FC=fold change.

## References

1. Fowler DM, Araya CL, Fleishman SJ, Kellogg EH, Stephany JJ, Baker D, et al. High-resolution mapping of protein sequence-function relationships. Nat Methods. 2010;7:741–6.

2. Starita LM, Young DL, Islam M, Kitzman JO, Gullingsrud J, Hause RJ, et al. Massively Parallel Functional Analysis of BRCA1 RING Domain Variants. Genetics. 2015;200:413–22.

3. Majithia AR, Tsuda B, Agostini M, Gnanapradeepan K, Rice R, Peloso G, et al. Prospective functional classification of all possible missense variants in PPARG. Nat Genet. 2016;48:1570–5.

4. Weile J, Sun S, Cote AG, Knapp J, Verby M, Mellor JC, et al. A framework for exhaustively mapping functional missense variants. Mol Syst Biol. 2017;13:957.

5. Matreyek KA, Starita LM, Stephany JJ, Martin B, Chiasson MA, Gray VE, et al. Multiplex assessment of protein variant abundance by massively parallel sequencing. Nat Genet. Nature Publishing Group; 2018;50:874–82.

6. Kircher M, Xiong C, Martin B, Schubach M, Inoue F, Bell RJA, et al. Saturation mutagenesis of twenty disease-associated regulatory elements at single base-pair resolution. Nat Commun. 2019;10:3583.

7. Chiasson MA, Rollins NJ, Stephany JJ, Sitko KA, Matreyek KA, Verby M, et al. Multiplexed measurement of variant abundance and activity reveals VKOR topology, active site and human variant impact. Elife [Internet]. 2020;9. Available from:http://dx.doi.org/10.7554/eLife.58026

8. Starr TN, Greaney AJ, Hilton SK, Ellis D, Crawford KHD, Dingens AS, et al. Deep Mutational Scanning of SARS-CoV-2 Receptor Binding Domain Reveals Constraints on Folding and ACE2 Binding. Cell. 2020;182:1295–310.e20.

9. Suiter CC, Moriyama T, Matreyek KA, Yang W, Scaletti ER, Nishii R, et al. Massively parallel variant characterization identifies NUDT15 alleles associated with thiopurine toxicity. Proc Natl Acad Sci U S A. 2020;117:5394–401.

10. Sun S, Weile J, Verby M, Wu Y, Wang Y, Cote AG, et al. A proactive genotype-to-patient-phenotype map for cystathionine beta-synthase. Genome Med. 2020;12:13.

11. Starita LM, Ahituv N, Dunham MJ, Kitzman JO, Roth FP, Seelig G, et al. Variant Interpretation: Functional Assays to the Rescue. Am J Hum Genet. 2017;101:315–25.

12. Weile J, Roth FP. Multiplexed assays of variant effects contribute to a growing genotype–phenotype atlas. Hum Genet. 2018;137:665–78.

13. Gelman H, Dines JN, Berg J, Berger AH, Brnich S, Hisama FM, et al. Recommendations for the collection and use of multiplexed functional data for clinical variant interpretation. Genome Med. 2019;11:85.

14. Bloom JD. Software for the analysis and visualization of deep mutational scanning data. BMC Bioinformatics. 2015;16:168.

15. Rubin AF, Gelman H, Lucas N, Bajjalieh SM, Papenfuss AT, Speed TP, et al. A statistical framework for analyzing deep mutational scanning data. Genome Biol. 2017;18:150.

16. Faure AJ, Schmiedel JM, Baeza-Centurion P, Lehner B. DiMSum: an error model and pipeline for analyzing deep mutational scanning data and diagnosing common experimental pathologies. Genome Biol. 2020;21:207.

17. Sandmann S, de Graaf AO, Karimi M, van der Reijden BA, Hellström-Lindberg E, Jansen JH, et al. Evaluating Variant Calling Tools for Non-Matched Next-Generation Sequencing Data. Sci Rep. 2017;7:43169.

18. Cibulskis K, Lawrence MS, Carter SL, Sivachenko A, Jaffe D, Sougnez C, et al. Sensitive detection of somatic point mutations in impure and heterogeneous cancer samples. Nat Biotechnol. 2013;31:213–9.

19. Xu C, Gu X, Padmanabhan R, Wu Z, Peng Q, DiCarlo J, et al. smCounter2: an accurate low-frequency variant caller for targeted sequencing data with unique molecular identifiers. Bioinformatics. 2019;35:1299–309.

20. Zheng Z, Liebers M, Zhelyazkova B, Cao Y, Panditi D, Lynch KD, et al. Anchored multiplex PCR for targeted next-generation sequencing. Nat Med. 2014;20:1479–84.

21. Waltari E, Jia M, Jiang CS, Lu H, Huang J, Fernandez C, et al. 5’ Rapid Amplification of cDNA Ends and Illumina MiSeq Reveals B Cell Receptor Features in Healthy Adults, Adults With Chronic HIV-1 Infection, Cord Blood, and Humanized Mice.Front Immunol. 2018;9:628.

22. Lin Y-H, Hung S-J, Chen Y-L, Lin C-H, Kung T-F, Yeh Y-C, et al. Dissecting efficiency of a 5’ rapid amplification of cDNA ends (5’-RACE) approach for profiling T-cell receptor beta repertoire. PLoS One. 2020;15:e0236366.

23. Ewing AD, Houlahan KE, Hu Y, Ellrott K, Caloian C, Yamaguchi TN, et al. Combining tumor genome simulation with crowdsourcing to benchmark somatic single-nucleotide-variant detection. Nat Methods. 2015;12:623–30.

24. Rubin AF, Lucas N, Bajjalieh SM, Papenfuss AT, Speed TP, Fowler DM. Enrich2: a statistical framework for analyzing deep mutational scanning data [Internet]. Cold Spring Harbor Laboratory. 2016 [cited 2021 Mar 9]. p. 075150. Available from: https://www.biorxiv.org/content/10.1101/075150v1.abstract

25. Ma X, Shao Y, Tian L, Flasch DA, Mulder HL, Edmonson MN, et al. Analysis of error profiles in deep next-generation sequencing data. Genome Biol. 2019;20:50.

26. Davis EM, Sun Y, Liu Y, Kolekar P, Shao Y, Szlachta K, et al. SequencErr: measuring and suppressing sequencer errors in next-generation sequencing data. Genome Biol. 2021;22:37.

27. Stoler N, Nekrutenko A. Sequencing error profiles of Illumina sequencing instruments. NAR Genom Bioinform. 2021;3:lqab019.

28. Chen S, Zhou Y, Chen Y, Huang T, Liao W, Xu Y, et al. Gencore: an efficient tool to generate consensus reads for error suppressing and duplicate removing of NGS data. BMC Bioinformatics. 2019;20:606.

29. Matreyek KA, Stephany JJ, Chiasson MA, Hasle N, Fowler DM. An improved platform for functional assessment of large protein libraries in mammalian cells. Nucleic Acids Res. 2020;48:e1.

30. Meier M, Janosik M, Kery V, Kraus JP, Burkhard P. Structure of human cystathionine beta-synthase: a unique pyridoxal 5’-phosphate-dependent heme protein. EMBO J. 2001;20:3910–6.

31. Miles EW, Kraus JP. Cystathionine β-Synthase: Structure, Function, Regulation, and Location of Homocystinuria-causing Mutations *. J Biol Chem. Elsevier; 2004;279:29871–4.

32. Ereño-Orbea J, Majtan T, Oyenarte I, Kraus JP, Martínez-Cruz LA. Structural basis of regulation and oligomerization of human cystathionine β-synthase, the central enzyme of transsulfuration. Proc Natl Acad Sci U S A. 2013;110:E3790–9.

33. Oliveriusová J, Kery V, Maclean KN, Kraus JP. Deletion Mutagenesis of Human Cystathionine β-Synthase: Impact on activity, oligomeric status, ands-adenosylmethionine regulation. Journal of Biological [Internet]. ASBMB; 2002; Available from:https://www.jbc.org/article/S0021-9258(19)33070-4/abstract

34. Majtan T, Singh LR, Wang L, Kruger WD. Active cystathionine β-synthase can be expressed in heme-free systems in the presence of metal-substituted porphyrins or a chemical chaperone. Journal of Biological [Internet]. ASBMB; 2008; Available from:https://www.jbc.org/article/S0021-9258(20)63280-X/abstract

35. Kozich V, Sokolová J, Klatovská V, Krijt J, Janosík M, Jelínek K, et al. Cystathionine beta-synthase mutations: effect of mutation topology on folding and activity. Hum Mutat. Wiley; 2010;31:809–19.

36. Majtan T, Liu L, Carpenter JF, Kraus JP. Rescue of Cystathionine β-Synthase (CBS) Mutants with Chemical Chaperones: PURIFICATION AND CHARACTERIZATION OF EIGHT CBS MUTANT ENZYMES *. J Biol Chem. Elsevier; 2010;285:15866–73.

37. Casique L, Kabil O, Banerjee R, Martinez JC, De Lucca M. Characterization of two pathogenic mutations in cystathionine beta-synthase: Different intracellular locations for wild-type and mutant proteins [Internet]. Gene. 2013. p. 117–24. Available from:http://dx.doi.org/10.1016/j.gene.2013.08.021

38. Kery V, Bukovska G, Kraus JP. Transsulfuration depends on heme in addition to pyridoxal 5’-phosphate. Cystathionine beta-synthase is a heme protein. J Biol Chem. 1994;269:25283–8.

39. Komar AA, Kommer A, Krasheninnikov IA, Spirin AS. Cotranslational heme binding to nascent globin chains. FEBS Lett. 1993;326:261–3.

40. Komar AA, Kommer A, Krasheninnikov IA, Spirin AS. Cotranslational folding of globin. J Biol Chem. 1997;272:10646–51.

41. Balchin D, Hayer-Hartl M, Hartl FU. In vivo aspects of protein folding and quality control. Science. 2016;353:aac4354.

42. Joazeiro CAP. Mechanisms and functions of ribosome-associated protein quality control. Nat Rev Mol Cell Biol. 2019;20:368–83.

43. Shih VE, Fringer JM, Mandell R, Kraus JP, Berry GT, Heidenreich RA, et al. A missense mutation (I278T) in the cystathionine beta-synthase gene prevalent in pyridoxine-responsive homocystinuria and associated with mild clinical phenotype. Am J Hum Genet. 1995;57:34–9.

44. de Franchis R, Kraus E, Kozich V, Sebastio G, Kraus JP. Four novel mutations in the cystathionine beta-synthase gene: effect of a second linked mutation on the severity of the homocystinuric phenotype. Hum Mutat. Wiley; 1999;13:453–7.

45. Gaustadnes M, Wilcken B, Oliveriusova J, McGill J, Fletcher J, Kraus JP, et al. The molecular basis of cystathionine beta-synthase deficiency in Australian patients: genotype-phenotype correlations and response to treatment. Hum Mutat. 2002;20:117–26.

46. Mendes MIS, Colaço HG, Smith DEC, Ramos RJJF, Pop A, van Dooren SJM, et al. Reduced response of Cystathionine Beta-Synthase (CBS) to S-Adenosylmethionine (SAM): Identification and functional analysis of CBS gene mutations in Homocystinuria patients. J Inherit Metab Dis. Wiley; 2014;37:245–54.

47. Presnyak V, Alhusaini N, Chen Y-H, Martin S, Morris N, Kline N, et al. Codon optimality is a major determinant of mRNA stability. Cell. 2015;160:1111–24.

48. Bazzini AA, del Viso F, Moreno-Mateos MA, Johnstone TG, Vejnar CE, Qin Y, et al. Codon identity regulates mRNA stability and translation efficiency during the maternal-to-zygotic transition. EMBO J. 2016;35:2087–103.

49. Gamble CE, Brule CE, Dean KM, Fields S, Grayhack EJ. Adjacent Codons Act in Concert to Modulate Translation Efficiency in Yeast. Cell. 2016;166:679–90.

50. Wu Q, Medina SG, Kushawah G, DeVore ML, Castellano LA, Hand JM, et al. Translation affects mRNA stability in a codon-dependent manner in human cells. Elife [Internet]. 2019;8. Available from:http://dx.doi.org/10.7554/eLife.45396

51. Narula A, Ellis J, Taliaferro JM, Rissland OS. Coding regions affect mRNA stability in human cells. RNA. 2019;25:1751–64.

52. Medina-Muñoz SG, Kushawah G, Castellano LA, Diez M, DeVore ML, Salazar MJB, et al. Crosstalk between codon optimality and cis-regulatory elements dictates mRNA stability. Genome Biol. 2021;22:14.

53. Behrens A, Rodschinka G, Nedialkova DD. High-resolution quantitative profiling of tRNA abundance and modification status in eukaryotes by mim-tRNAseq. Mol Cell. 2021;81:1802–15.e7.

54. Gogakos T, Brown M, Garzia A, Meyer C, Hafner M, Tuschl T. Characterizing Expression and Processing of Precursor and Mature Human tRNAs by Hydro-tRNAseq and PAR-CLIP [Internet]. Cell Reports. 2017. p. 1463–75. Available from:http://dx.doi.org/10.1016/j.celrep.2017.07.029

55. Taoka S, Lepore BW, Kabil Ö, Ojha S, Ringe D, Banerjee R. Human cystathionine β-synthase is a heme sensor protein. Evidence that the redox sensor is heme and not the vicinal cysteines in the CXXC motif seen in the crystal structure of the truncated enzyme. Biochemistry. American Chemical Society (ACS); 2002;41:10454–61.

56. Tewhey R, Kotliar D, Park DS, Liu B, Winnicki S, Reilly SK, et al. Direct Identification of Hundreds of Expression-Modulating Variants using a Multiplexed Reporter Assay [Internet]. Cell. 2018. p. 1132–4. Available from:http://dx.doi.org/10.1016/j.cell.2018.02.021

57. Sample PJ, Wang B, Reid DW, Presnyak V, McFadyen IJ, Morris DR, et al. Human 5’ UTR design and variant effect prediction from a massively parallel translation assay. Nat Biotechnol. 2019;37:803–9.

58. Lim Y, Arora S, Schuster SL, Corey L, Fitzgibbon M, Wladyka CL, et al. Multiplexed functional genomic analysis of 5’untranslated region mutations across the spectrum of prostate cancer. Nat Commun. Nature Publishing Group; 2021;12:1–18.

59. Nainar S, Cuthbert BJ, Lim NM, England WE, Ke K, Sophal K, et al. An optimized chemical-genetic method for cell-specific metabolic labeling of RNA. Nat Methods. 2020;17:311–8.

60. Findlay GM, Daza RM, Martin B, Zhang MD, Leith AP, Gasperini M, et al. Accurate classification of BRCA1 variants with saturation genome editing. Nature. 2018;562:217–22.

61. Duan J, Wainwright MS, Comeron JM, Saitou N, Sanders AR, Gelernter J, et al. Synonymous mutations in the human dopamine receptor D2 (DRD2) affect mRNA stability and synthesis of the receptor. Hum Mol Genet. 2003;12:205–16.

62. Nackley AG, Shabalina SA, Tchivileva IE, Satterfield K, Korchynskyi O, Makarov SS, et al. Human catechol-O-methyltransferase haplotypes modulate protein expression by altering mRNA secondary structure. Science. 2006;314:1930–3.

63. Duan J, Shi J, Ge X, Dölken L, Moy W, He D, et al. Genome-wide survey of interindividual differences of RNA stability in human lymphoblastoid cell lines. Sci Rep. 2013;3:1318.

64. Li Q, Makri A, Lu Y, Marchand L, Grabs R, Rousseau M, et al. Genome-wide search for exonic variants affecting translational efficiency. Nat Commun. 2013;4:2260.

65. Battle A, Khan Z, Wang SH, Mitrano A, Ford MJ, Pritchard JK, et al. Genomic variation. Impact of regulatory variation from RNA to protein. Science. 2015;347:664–7.

66. Cenik C, Cenik ES, Byeon GW, Grubert F, Candille SI, Spacek D, et al. Integrative analysis of RNA, translation, and protein levels reveals distinct regulatory variation across humans. Genome Res. 2015;25:1610–21.

67. Kirchner S, Cai Z, Rauscher R, Kastelic N, Anding M, Czech A, et al. Alteration of protein function by a silent polymorphism linked to tRNA abundance. PLoS Biol. 2017;15:e2000779.

68. Zhou T, Weems M, Wilke CO. Translationally optimal codons associate with structurally sensitive sites in proteins. Mol Biol Evol. 2009;26:1571–80.

69. Li H, Handsaker B, Wysoker A, Fennell T, Ruan J, Homer N, et al. The Sequence Alignment/Map format and SAMtools. Bioinformatics. 2009;25:2078–9.

70. Heger A, Belgrad TG, Goodson M, Jacobs K. pysam: Python interface for the SAM/BAM sequence alignment and mapping format. 2014.

71. Langmead B, Trapnell C, Pop M, Salzberg SL. Ultrafast and memory-efficient alignment of short DNA sequences to the human genome. Genome Biol. 2009;10:R25.

72. Martin M. Cutadapt removes adapter sequences from high-throughput sequencing reads. EMBnet.journal. 2011;17:10–2.

73. Smith T, Heger A, Sudbery I. UMI-tools: modeling sequencing errors in Unique Molecular Identifiers to improve quantification accuracy. Genome Res. 2017;27:491–9.

74. Dowle M, Srinivasan A, Gorecki J, Chirico M, Stetsenko P, Short T, et al. Package “data. table.” Extension of ‘data frame [Internet]. 2019; Available from: ftp://ftp.musicbrainz.org/pub/cran/web/packages/data.table/data.table.pdf

75. Wickham H. ggplot2. WIREs Comp Stat. 2011;3:180–5.

76. Wilke CO. >cowplot: streamlined plot theme and plot annotations for “ggplot2.” CRAN Repos. 2016;2:R2.

77. Garnier, Simon, Ross, Noam, Rudis, Robert, et al. viridis - Colorblind-Friendly Color Maps for R [Internet]. 2021. Available from: https://sjmgarnier.github.io/viridis/

78. Xiao N. ggsci: scientific journal and sci-fi themed color palettes for “ggplot2.” R package version. 2018;2.

79. Lumley T, Lumley MT. Package “leaps.” Regression subset selection Thomas Lumley Based on Fortran Code by Alan Miller Available online: http://CRANR-projectorg/package=leaps (Accessed on 18 March 2018) [Internet]. 2013; Available from: https://cran.microsoft.com/snapshot/2016-08-29/web/packages/leaps/leaps.pdf

80. Kuhn M, Wing J, Weston S, Williams A, Keefer C, Engelhardt A, et al. Package “caret.” R J. 2020;223:7.

81. Meyer D, Dimitriadou E, Hornik K, Weingessel A, Leisch F, Chang C-C, et al. Package “e1071.” R J [Internet]. 2019; Available from: http://sunsite2.icm.edu.pl/pub/unix/math/cran/web/packages/e1071/e1071.pdf

82. Venables WNRipley BD. Modern applied statistics with S. Statistics and computing New York: Springer. 2002;

83. Ripley BD. Pattern Recognition and Neural Networks. Cambridge University Press; 2007.

84. Liaw A, Wiener M, Others. Classification and regression by randomForest. R news. 2002;2:18–22.

85. Ridgeway G, Ridgeway MG. The gbm package. R Foundation for Statistical Computing, Vienna, Austria [Internet]. 2004;5. Available from:https://ftp.uni-bayreuth.de/math/statlib/R/CRAN/doc/packages/gbm.pdf

86. Friedman J, Hastie T, Tibshirani R, Narasimhan B. Package “glmnet.” CRAN R Repositary [Internet]. 2021; Available from: http://masterdistfiles.gentoo.org/pub/cran/web/packages/glmnet/glmnet.pdf

87. Smyth GK. limma: Linear Models for Microarray Data. In: Gentleman R, Carey VJ, Huber W, Irizarry RA, Dudoit S, editors. Bioinformatics and Computational Biology Solutions Using R and Bioconductor. New York, NY: Springer New York; 2005. p. 397–420.

