## Additional File 1 for "satmut_utils: a simulation and variant calling package for multiplexed assays of variant effect"

Variant conversion

We coined the term *variant conversion* for cases where a true positive variant is phased with a nearby true or false positive variant, leading to miscalls. We have observed significant variant conversion when the variant calling algorithm is not constrained in the merging of nearby mismatches into primary MNP variant calls (see Considerations for variant calling algorithm design). Erroneous variant calls due to conversion are dependent on the product of 1) the probability of a true variant occurring at a given position, and 2) the probability of error in a window surrounding the position. For satmut_utils, this window is 3 nt. However, other variant callers may not impose any window constraint when merging mismatches into MNP; in this case, variant conversion can be a significant source of both false positives and false negatives.

Variant frequencies in saturation mutagenesis libraries

Variant frequencies are primarily dictated by the number of species in the variant library. As the size of the mutagenized target region increases, frequencies are driven lower as each variant-containing read contributes to depth at other target positions covered by the read. We recommend generating variant libraries that do not exceed approximately 5000 species, so that SNP variant frequencies are appreciably higher than typical PCR error rates (1 x 10^-7^ to 1 x 10^-4^) and so that libraries do not need to be sequenced with coverage depths greater than ~1 x 10^6^ per target position. Another consideration is that MNPs are generally represented at 10-fold lower frequency than SNPs in variant libraries, due to lower mutagenesis efficiency by virtue of unfavorable thermodynamics of degenerate primer annealing. Nonetheless, MNPs are more likely to represent true variants over SNPs, as the probability of observing two and three mismatches to the reference is significantly less than the probability of observing a single mismatch in near reference-space. Thus, when designing experiments, one should consider not only the number of variants to mutagenize, but also the relative composition of SNPs and MNPs in the library.

Comment on existing simulation tools

Several solutions exist for simulating sequencing reads with variants [[1–4]](https://paperpile.com/c/B4R8cI/rwSkU+UmP8w+rK2lW+B2dDQ). These have successfully been used for simulating cancer genomes to assess germline and somatic variant calling tools [[5–11]](https://paperpile.com/c/B4R8cI/zDHmt+YAAxi+oNV54+8ziBU+duVsV+ss4R3+7hrFr). Most of the simulators learn an error model from real alignments and then generate reads and variants starting from a reference sequence. In contrast, BAMSurgeon [[12]](https://paperpile.com/c/B4R8cI/q7LSX) and satmut_utils edit variants into real alignments. This approach captures the native error profile attributable to the library preparation and sequencing methods.

Simulation algorithm design

Using read sampling and a heuristic, satmut_utils ‘sim’ addresses the problem of simulation of many low-frequency variants at the same (and nearby) positions, while enforcing that no read pair is edited more than once. To avoid variant conversion, ‘sim’ provides the –edit_buffer argument to enable the user to only simulate variants in read pairs when the read matches the reference sequence +/- the edit buffer about the variant coordinate(s). This allows the user to control how “idealized” simulation is, which may affect interpretation of downstream variant calling performance. For example, if a low-frequency variant is simulated with only one count, and the randomly selected read pair to edit has an error nearby within the edit buffer, the variant may be not be called (depending on the variant calling algorithm). In this case, increasing the edit buffer ensures the variant is simulated in a read pair without creating a higher order variant (e.g. SNP to a di-nt MNP composed of the true SNP and one error). Similar control of which reads are edited is also provided by the –max_nm argument. If at least one read of a pair has an edit distance greater than the max edit distance, the pair is disqualified for editing.

Simulation requirements for assessing performance of consensus deduplication

The satmut_utils ‘sim’ workflow currently does not meet the requirements to measure the sensitivity-specificity tradeoff of consensus deduplication. To do this, reads must be grouped by unique molecular index (UMI) prior to simulation, and then the ‘sim’ workflow would need to make use of this information to ensure that all duplicates within a UMI group are edited. (Otherwise, the variant is not retained in the consensus). In summary, the requirements to properly simulate variants to measure the improvement of consensus deduplication pose significant challenges that preclude its current support. The use of biological standards of known variant composition are better suited to measure the performance improvement of deduplication.

Considerations for variant calling algorithm design

A primary consideration for the algorithmic design of a MAVE variant caller is how to implement MNP calling. Specifically, should MNP calls be made across the entire read or confined to a smaller window? The downside of imposing no window constraint is that mismatches spanning greater than a small window are more likely to be false positives due to variant conversion. When simulating MNPs with large spans (haplotypes) and calling variants with a prototype algorithm that does not require mismatches to be within a span/window of 3 nt, we observed that the number of false positive variants scaled rapidly as the window size increased. Even with a relatively short window of 10 nt, the number of false positive variants exploded due to variant conversion and merging of errors.

Because mutagenesis primer lengths are almost always >10 nt, it is unexpected to observe phased variants shorter than a typical primer length (20 nt) if mutagenesis is carried out by a single primer extension reaction with pooled mutagenesis oligos [[13]](https://paperpile.com/c/B4R8cI/VN3od). Thus, we opted to only merge mismatches within a 3 nt window, which implies true haplotypes with mismatches spanning 4 nt or greater will not be called together by satmut_utils. We conclude that without improved variant calling algorithms and/or error correction methods, long-range, low-frequency haplotypes cannot be reliably called without incurring a high cost of false positives.

Finally, while InDel (insertion and deletion) rates may be predictive of library preparation or sequencing failure, the satmut_utils workflow does not filter out reads with InDels, as do dms_tools2 and Enrich2. That is, variant calls may still be made from a read pair even if there is a small InDel somewhere else in the read(s).

Benchmarking considerations

We expect that the lack of benchmarking from prior MAVE variant callers is due to difficulty of configuration and specific constraints on input data. For example, to successfully benchmark against Enrich2 and dms_tools2, we wrote a script to 1) filter reads containing InDels, 2) make reads flush with codons by trimming and/or appending reference sequence, and 3) add barcodes/UMIs to the 5’ end of each read. We used the same filtered alignments for simulation and benchmarking of all variant callers, with the exception that we added 12 nt barcodes to each read for dms_tools2. Although an alignment strategy (DiMSum, satmut_utils) is more computationally expensive, it imposes no constraints on primer design or library preparation chemistry.

One explanation for lower precision of Enrich2 at perfect recall was our use of its Basic mode, which does not consider base call support from both reads. We attempted to use Enrich2 Overlap mode, but we found a high proportion of variant calls were unresolvable (<https://github.com/FowlerLab/Enrich2/issues/45>), precluding complete analysis of all simulated variants. Another reason Enrich2 and DimSum may have poorer precision than satmut_utils is that the former variant calling methods may not put a constraint on the merging of mismatches into MNP calls (see Considerations for variant calling algorithm design).

Lastly, one other possible explanation for differences in performance is that analysis parameters for read filtering (e.g. base quality threshold) are implemented in different ways for each variant caller. Nonetheless, in benchmarking, efforts were made to make analysis parameters as similar as possible between variant callers. Also, by simulating variants with a NNK mutagenesis signature, we benchmarked the improvement in precision possible by filtering on the mutagenesis signature. The satmut_utils ‘call’ workflow annotates the signature for each variant and enables the exclusion of variants that do not match it, which partially accounts for the boost in precision for satmut_utils relative to other variant callers.

Time and memory consumption

6,359,057 read pairs from two amplicons analyzed with satmut_utils ‘call’ (with primer base quality masking, –nthreads 10) took 42 min and consumed a maximum of 142.2 Mb memory.

42,900,903 read pairs from fifteen tiles (RACE-like chemistry) analyzed with satmut_utils ‘call’ (with primer base quality masking and consensus deduplication, –nthreads 10) took 16 h and consumed a maximum of 165 Mb memory.

We have successfully analyzed up to 67 million read pairs (RACE-like chemistry, primer base quality masking, consensus deduplication, –nthreads 10), which took nearly 47 h. Consensus deduplication can take significant time and will be optimized in future versions.

Error correction model filtering

satmut_utils call extracts numerous quality features for the mismatches comprising both SNP and MNP variant calls. That is, for MNPs, quality features are provided for each mismatch in a primary variant call. This allows filtering of MNPs composed of a true variant and an error, which should help remediate variant conversion. After application of ML models for filtering individual mismatches, the user must appropriately filter MNP variants with both true and false positive predictions. That is, if a component mismatch of a MNP call is determined to be false by the error correction model, the user must manually filter the encompassing variant call, using prior knowledge of the original variant type (di-nt MNP, tri-nt MNP) and how many mismatches were retained in the variant call after filtering.

Caveats of error correction models

While our results suggest binary classifiers can learn error signatures in each experiment, there is a risk for bias when applying models to real data. Models trained on simulated data are unlikely to generalize on real datasets if the ‘sim’ parameters (variant frequencies, SNP/MNP composition) are not representative of the true probability distributions for real mutagenesis libraries. Furthermore, regarding the high performance of error correction models, we caution that the design rules of satmut_utils ‘sim’ (see Simulation algorithm design) impose constraints on read editing (mate pair overlap, edit distance, edit buffer), that may be subsequently learned by downstream models. We also did not include ‘sim’ hyperparameters under the scope of cross-validation. Together, these limitations may lead to optimistic performance measures [[14]](https://paperpile.com/c/B4R8cI/Ub3MI).

References

1. Huang W, Li L, Myers JR, Marth GT. ART: a next-generation sequencing read simulator. Bioinformatics. 2012;28:593–4.

2. Pattnaik S, Gupta S, Rao AA, Panda B. SInC: an accurate and fast error-model based simulator for SNPs, Indels and CNVs coupled with a read generator for short-read sequence data. BMC Bioinformatics. 2014;15:40.

3. Mu JC, Mohiyuddin M, Li J, Bani Asadi N, Gerstein MB, Abyzov A, et al. VarSim: a high-fidelity simulation and validation framework for high-throughput genome sequencing with cancer applications. Bioinformatics. 2015;31:1469–71.

4. Stephens ZD, Hudson ME, Mainzer LS, Taschuk M, Weber MR, Iyer RK. Simulating Next-Generation Sequencing Datasets from Empirical Mutation and Sequencing Models. PLoS One. 2016;11:e0167047.

5. Li H. A statistical framework for SNP calling, mutation discovery, association mapping and population genetical parameter estimation from sequencing data. Bioinformatics. 2011;27:2987–93.

6. Larson DE, Harris CC, Chen K, Koboldt DC, Abbott TE, Dooling DJ, et al. SomaticSniper: identification of somatic point mutations in whole genome sequencing data. Bioinformatics. 2012;28:311–7.

7. Cibulskis K, Lawrence MS, Carter SL, Sivachenko A, Jaffe D, Sougnez C, et al. Sensitive detection of somatic point mutations in impure and heterogeneous cancer samples. Nat Biotechnol. 2013;31:213–9.

8. do Valle ÍF, Giampieri E, Simonetti G, Padella A, Manfrini M, Ferrari A, et al. Optimized pipeline of MuTect and GATK tools to improve the detection of somatic single nucleotide polymorphisms in whole-exome sequencing data. BMC Bioinformatics. 2016;17:341.

9. Kim S, Scheffler K, Halpern AL, Bekritsky MA, Noh E, Källberg M, et al. Strelka2: fast and accurate calling of germline and somatic variants. Nat Methods. 2018;15:591–4.

10. Narzisi G, Corvelo A, Arora K, Bergmann EA, Shah M, Musunuri R, et al. Genome-wide somatic variant calling using localized colored de Bruijn graphs. Commun Biol. 2018;1:20.

11. Xu C, Gu X, Padmanabhan R, Wu Z, Peng Q, DiCarlo J, et al. smCounter2: an accurate low-frequency variant caller for targeted sequencing data with unique molecular identifiers. Bioinformatics. 2019;35:1299–309.

12. Ewing AD, Houlahan KE, Hu Y, Ellrott K, Caloian C, Yamaguchi TN, et al. Combining tumor genome simulation with crowdsourcing to benchmark somatic single-nucleotide-variant detection. Nat Methods. 2015;12:623–30.

13. Weile J, Sun S, Cote AG, Knapp J, Verby M, Mellor JC, et al. A framework for exhaustively mapping functional missense variants. Mol Syst Biol. 2017;13:957.

14. Cawley GC, Talbot NLC. On over-fitting in model selection and subsequent selection bias in performance evaluation. J Mach Learn Res. JMLR. org; 2010;11:2079–107.
